# Mitochondrial genome instability disrupts brown adipose tissue through pseudohypoxia-iron-NAD⁺ axis

**DOI:** 10.64898/2026.08.28.747463

**Authors:** Serena S Özturk, Swagat Pradhan, Madeleine H Lackman, Lipsa Rani Panda, Aleksandra Zhaivoron, Markus Innila, João Santos Patricio, Lauren Zacharias, Thomas Mathews, Sinem Karaman, Nahid A Khan

## Abstract

Brown adipose tissue (BAT) is a mitochondria-rich thermogenic organ whose function depends on high oxidative capacity, yet how primary mitochondrial dysfunction remodels BAT identity and metabolism remains poorly defined. Using the Deletor mouse model of progressive mtDNA deletion disease, we identify a pseudohypoxia-iron-NAD^+^ axis as a central organiser of BAT pathology. Deletor BAT underwent profound structural, transcriptional and metabolic remodelling, characterised by mitochondrial ultrastructural damage, loss of thermogenic identity, PHD3/HIF-associated pseudohypoxic signalling, iron dysregulation and NAD^+^/NADH redox imbalance. Indirect calorimetry confirmed that this molecular disease program translates to functional thermogenic failure under physiological demand. Whole-body indirect calorimetry showed reduced oxygen consumption during acute cold challenge and impaired RER-based fuel switching, consistent with impaired cold-adaptive oxidative metabolism. Metabolomic profiling revealed altered TCA cycle intermediates, glycolytic rewiring and selective amino acid accumulation. Pharmacological perturbation showed that the PHD inhibitor roxadustat worsened disease-associated features, whereas HIF-1α suppression with PX-478 attenuated the integrated stress response, indicating that pseudohypoxic signalling is maladaptive in this setting. Nicotinamide riboside broadly attenuated the disease metabolome and transcriptome, restoring NAD^+^/NADH balance, suppressing ISRmt, iron-stress and pseudohypoxic gene programs, and correcting selective carnitine and acylcarnitine abnormalities consistent with impaired fatty-acid handling. These findings define a therapeutically tractable pseudohypoxia-iron-NAD^+^ axis as a core determinant of BAT dysfunction in mitochondrial disease.

## Introduction

Brown adipose tissue (BAT) is mitochondria-enriched metabolic organ specialized for adaptive thermogenesis and systemic energy homeostasis. Unlike white adipose tissue, which stores excess energy primarily in the form of triglycerides, BAT dissipates chemical energy as heat through mitochondrial respiration and uncoupling protein 1 (UCP1), thereby contributing to whole-body glucose, lipid, and redox balance while also communicating with peripheral tissues through secreted metabolic signals [1–4]. This thermogenic function depends on high mitochondrial density and sustained respiratory chain activity, not only for ATP generation and substrate oxidation but also for TCA cycle, redox balance, and fatty acid handling [5,6], making BAT potentially vulnerable to defects in mitochondrial genome maintenance and respiratory chain function. Mitochondrial diseases caused by instability of mitochondrial DNA (mtDNA) provide a powerful framework in which to address this question. The Deletor mouse, which accumulates multiple mtDNA deletions due to expression of a dominant patient mutation in Twinkle helicase, develops progressive respiratory chain deficiency and recapitulates key features of human mitochondrial myopathy caused by mtDNA maintenance defects[7,8]. Most work in this model has focused on skeletal muscle, where mitochondrial ultrastructural abnormalities, respiratory chain failure, and integrated stress responses drive progressive muscle phenotype [9–13]. Brown adipocytes require intact respiratory chain function not only for ATP generation and substrate oxidation, but also for tricarboxylic acid (TCA) cycle activity, redox balance, fatty acid handling, and maintenance of thermogenic identity [14,15]. Whether chronic mtDNA instability induces a distinct BAT specific disease remodelling, and which pathways govern its adaptation or compensation, remain unknown.

The oxygen-sensing prolyl hydroxylase-hypoxia-inducible factor (PHD-HIF) pathway provides a strong mechanistic framework for understanding BAT remodelling under mitochondrial stress. Under normoxic conditions, PHD enzymes hydroxylate HIF-α subunits and target them for degradation, whereas respiratory failure or altered intracellular metabolite availability can stabilize HIF signalling despite normal environmental oxygen tension, producing pseudohypoxia [16–18]. Among the canonical PHD isoforms, PHD3/*Egln3* is highly stress-responsive and closely coupled to cellular metabolic state, because its activity depends on α-ketoglutarate, ferrous iron, oxygen and redox balance, all of which are perturbed by mitochondrial dysfunction [19–21]. While HIF signalling in adipose tissue has been studied largely in obesity models and PHD2 biology [22] its role in BAT remodelling during primary mitochondrial dysfunction remains essentially unexplored. This oxygen-sensing axis does not act in isolation: PHD3 activity is itself a readout of iron availability, linking pseudohypoxia directly to iron metabolism, on which BAT is uniquely dependent [23–26]. Iron is essential for iron-sulphur cluster biogenesis, heme synthesis, electron transport chain assembly, and the function of numerous mitochondrial enzymes, yet excess labile iron drives oxidative stress and tissue damage positioning iron handling as both a cofactor for PHD3 and a liability for mitochondrial integrity. Disturbances in iron and heme metabolism could therefore simultaneously impair respiration, disrupt substrate catabolism, and amplify cellular stress [27]. A third, tightly linked layer of regulation is NAD^+^ metabolism. Because NAD^+^/NADH cycling is inseparable from electron transport and substrate oxidation, any disruption to the respiratory chain necessarily reverberates through cellular redox state. Our group previously identified NAD^+^ deficiency as a hallmark of mitochondrial myopathy [28,29], raising the question of whether analogous dysregulation contributes to BAT pathology. Whether mitochondrial dysfunction in BAT converges on a coordinated pseudohypoxia-iron-NAD^+^ axis, and whether this state is pharmacologically modulated, has not been defined.

Here, we show that Deletor BAT undergoes profound structural, transcriptional, and metabolic remodelling, characterised by loss of thermogenic identity, mitochondrial ultrastructural damage, activation of a pseudohypoxia-associated program centered on PHD3/HIF signalling, dysregulation of iron metabolism pathways, and NAD^+^ redox imbalance. Bidirectional pharmacological modulation of this axis provided functional evidence that this pseudohypoxic program is maladaptive: HIF stabilisation with roxadustat exacerbated BAT pathology, whereas HIF suppression with PX-478 attenuated the mitochondrial integrated stress response, demonstrating that the pseudohypoxic program is maladaptive rather than compensatory in this context. Iron loading independently aggravated fibrotic and metabolic stress, while nicotinamide riboside restored NAD^+^ redox balance and broadly normalized the BAT metabolome. Together, these findings identify pseudohypoxia-iron-NAD^+^ axis as a central organiser of BAT pathology in mitochondrial disease and provides a new framework for therapeutic intervention.

## Results

### mtDNA instability disrupts brown adipose tissue architecture and thermogenic identity

Brown adipose tissue depends on continuous mitochondrial oxidative capacity to support thermogenesis, making it potentially vulnerable to primary defects in mitochondrial genome maintenance. To determine whether chronic mtDNA instability remodels BAT structure and metabolism, we performed histological, ultrastructural, transcriptomic, and metabolomic analyses of interscapular BAT from Deletor mice, a progressive model of mitochondrial myopathy driven by accumulation of multiple mtDNA deletions and age-matched wild-type controls.

Electron microscopy revealed marked mitochondrial abnormalities in Deletor BAT, including severe disruption of cristae architecture, electron-lucent matrix spaces, and aberrant mitochondrial morphology (Fig. 1A). Cristae density was significantly reduced relative to wild-type BAT, demonstrating impairment of the mitochondrial ultrastructure that underpins thermogenic function (Fig. 1C). Consistent with the presence of mitochondrial genome instability in this tissue, long-range PCR detected multiple mtDNA deletion products in Deletor BAT (Supp. Fig. 1A). Oil Red O staining further showed that Deletor BAT has smaller and less abundant lipid droplets than wild-type BAT (Fig. 1B), with significant reductions in both lipid droplet area and mean lipid droplet diameter (Fig. 1D) consistent with loss of the lipid-rich morphology characteristic of metabolically competent brown adipocytes. Immunostaining for SDHA, a nuclear-encoded subunit of mitochondrial Complex II revealed no significant change in SDHA abundance in Deletor BAT (Supp. Fig. 1B, C), suggesting that mitochondrial mass is maintained despite severe ultrastructural changes.

**Figure 1.**
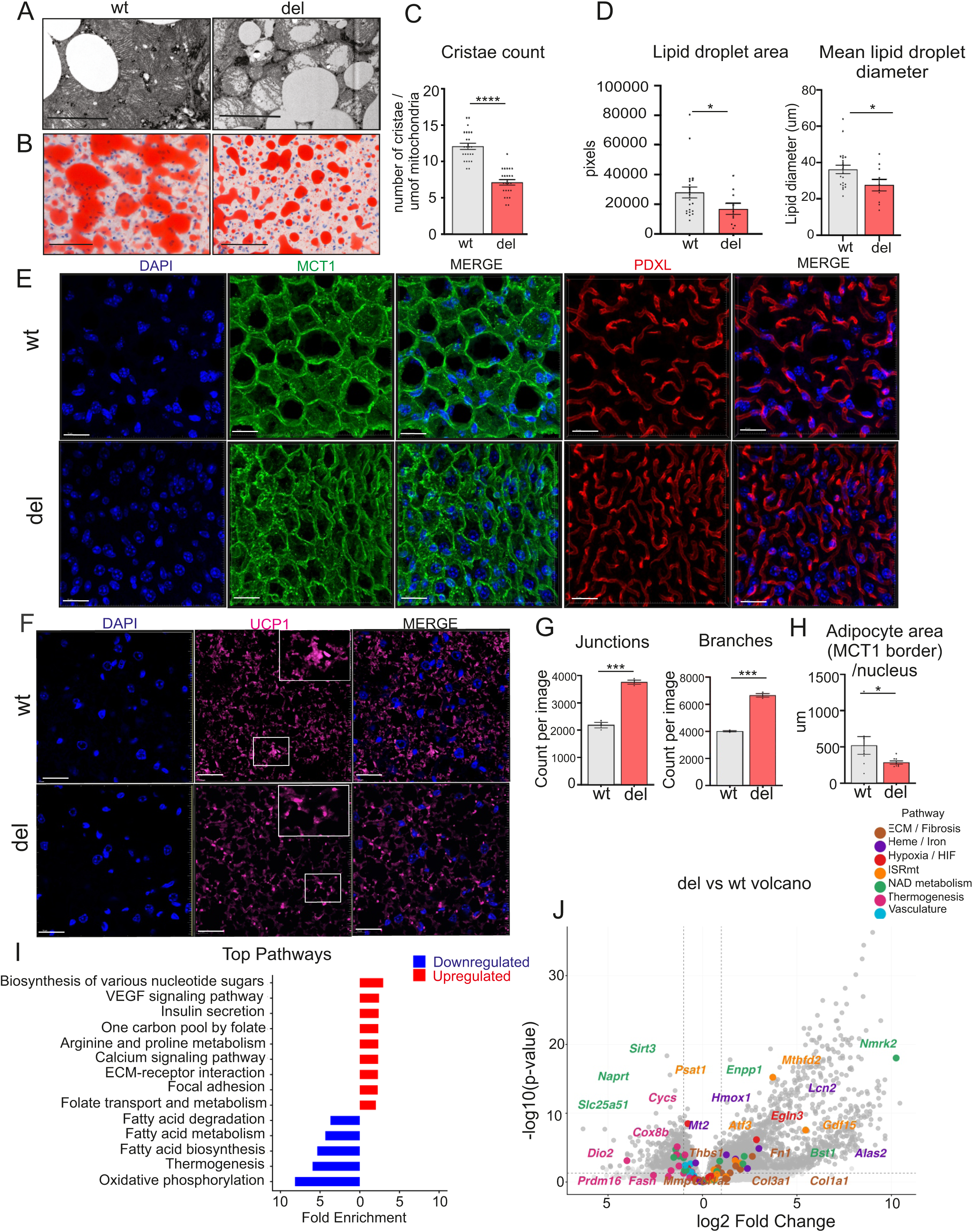
Mitochondrial dysfunction drives structural, and vascular remodelling in Deletor BAT. (A) Representative transmission electron micrographs of interscapular brown adipose tissue (BAT) from wild-type (WT) and Deletor (Del) mice showing abnormal mitochondrial morphology in Deletor BAT, including disrupted cristae architecture and electron-lucent matrix spaces. Scale bar =1µm (B) Representative Oil Red O staining of BAT sections from wild-type (WT) and Deletor (Del) mice showing reduced lipid droplet abundance and size in Deletor BAT. Scale bar = 200 µm (C) Quantification of mitochondrial cristae number per um of mitochondrial length in wild-type (WT) and Deletor (Del) BAT (n=24 per group). (D) Quantification of lipid droplet size and mean lipid droplet diameter in wild-type (WT) and Deletor (Del) BAT (n=21) WT (n=10) Del. (E) Representative immunofluorescence images of BAT sections stained for DAPI (blue), MCT1 (green), and podocalyxin (PDXL; red). MCT1 outlines brown adipocyte boundaries, and PDXL marks the vascular endothelium. Scale bar = 20 µm. (F) Representative immunofluorescence images of BAT sections stained for DAPI (blue) and UCP1 (magenta), showing reduced UCP1 immunoreactivity in Deletor BAT. Scale bar = 20 µm. (G) Quantification of vascular junctions count and vascular branches count per image in wild-type (WT) and Deletor (Del) (n=3 per group). (H) Quantification of MCT1-defined cell area per nucleus (n=8 per group) in wild-type (WT) and Deletor (Del) BAT. (I) KEGG pathway enrichment analysis of significant differentially expressed genes in Deletor (Del) versus wild type (WT) BAT, showing the top significantly enriched upregulated and downregulated pathways. (J) Volcano plot showing the fold-change and statistical significance of the transcriptomic changes between wild-type (WT) and Deletor (Del) BAT, with selected genes colour-coded by pathway class. (n=5 (WT) and n = 4 (Del)) Data are shown as mean ± SEM with individual data points overlaid where applicable. Statistical significance was assessed by unpaired Student’s t-test. *p<0.05, **p<0.01, ***p<0.001, ****p<0.0001. For image quantification, 5 images were acquired from each animal and average was used for the analysis.

Monocarboxylate Transporter 1 (MCT1) immunostaining, which delineates brown adipocyte boundaries, together with podocalyxin (PDXL) immunostaining marking the vascular endothelium, revealed coordinated remodelling of both the cellular and vascular compartments in Deletor BAT (Fig. 1E). Deletor BAT showed reduced MCT1-defined cellular area per nucleus, consistent with altered adipocyte morphology and reduced lipid storage capacity (Fig 1H). In parallel, podocalyxin staining showed a denser and more disorganised capillary network, with a fine mesh-like branching pattern replacing the regular vascular architecture observed in wild-type tissue (Fig. 1E). Quantification confirmed significantly increased branch count and junction density, indicating angiogenic remodelling (Fig. 1G). Loss of BAT thermogenic identity was confirmed by UCP1 immunostaining. UCP1 immunoreactivity was markedly reduced throughout the BAT parenchyma (Fig. 1F).

To determine whether the molecular remodelling of Deletor BAT was accompanied by impaired cold-adaptive metabolism, whole-body metabolic parameters were assessed by indirect calorimetry during the light phase, dark phase, 12 hours each and acute cold challenge at 4°C for 6 hours. Oxygen consumption showed a consistent trend toward reduction in Deletor mice during the light and dark phases and was significantly attenuated during cold challenge, indicating impaired cold-induced oxidative metabolism (Supp Fig 1D). Together, these findings show that chronic mtDNA instability drives coordinated structural decompensation across multiple levels of BAT organisation: mitochondrial ultrastructure, lipid storage, adipocyte morphology, vascular architecture, and thermogenic identity. This establishes BAT as a metabolically specialized tissue that is highly vulnerable to primary mitochondrial genome maintenance defects.

### mtDNA instability induces a coordinated pseudohypoxic and iron-stress disease program in Deletor BAT

To define the transcriptional landscape underlying these structural changes, we performed bulk RNA sequencing of Deletor and wild-type BAT. Differential expression analysis identified 3060 upregulated and 1988 downregulated out of 24044 genes in Deletor BAT relative to wild-type controls (Fig. 1J). Pathway annotation of differentially expressed genes in volcano plot revealed that the most significantly upregulated genes belonged to ISRmt, ECM/fibrosis, iron and heme metabolism, and HIF/hypoxia pathways, while thermogenesis and fatty acid metabolism genes were prominently downregulated and vascular remodelling genes were altered. KEGG pathway enrichment analysis confirmed that the most significantly upregulated pathways included VEGF signalling, arginine and proline metabolism, ECM-receptor interaction, one-carbon pool by folate, and focal adhesion, while the most significantly downregulated pathways were oxidative phosphorylation, thermogenesis, fatty acid biosynthesis, and fatty acid metabolism (Fig. 1I). Together, these transcriptional changes establish that structural remodelling in Deletor BAT is accompanied by a profound and coordinated shift in gene expression away from thermogenic and oxidative metabolism and toward integrated stress, vascular remodelling, and matrix remodelling programs.

Deletor BAT showed markedly elevated expression of integrated stress response genes *Mthfd2, Gdf15, Fgf21, Atf3, Atf4, Trib3, Psat1, and Phgdh* (Fig. 2A), consistent with broad activation of the ATF4/ATF5 linked stress axis and its downstream serine biosynthesis and one-carbon metabolic programs. Deletor BAT additionally showed increased expression of vascular remodelling genes, or genes associated with endothelial identity and angiogenic signalling, including *Angpt2* and *Nos3* (Fig. 2B), consistent with the angiogenic remodelling observed by podocalyxin immunostaining in Figure 1E, and indicating that vascular expansion is sustained by an active transcriptional program. Thermogenesis genes were coordinately suppressed, *Ucp1, Ppargc1a, Dio2*, and *Cidea* were all reduced relative to wild-type controls (Fig. 2D) corroborating the UCP1 protein loss observed by immunostaining and indicating that loss of thermogenic identity operates at the transcriptional and protein level in Deletor BAT.

**Figure 2.**
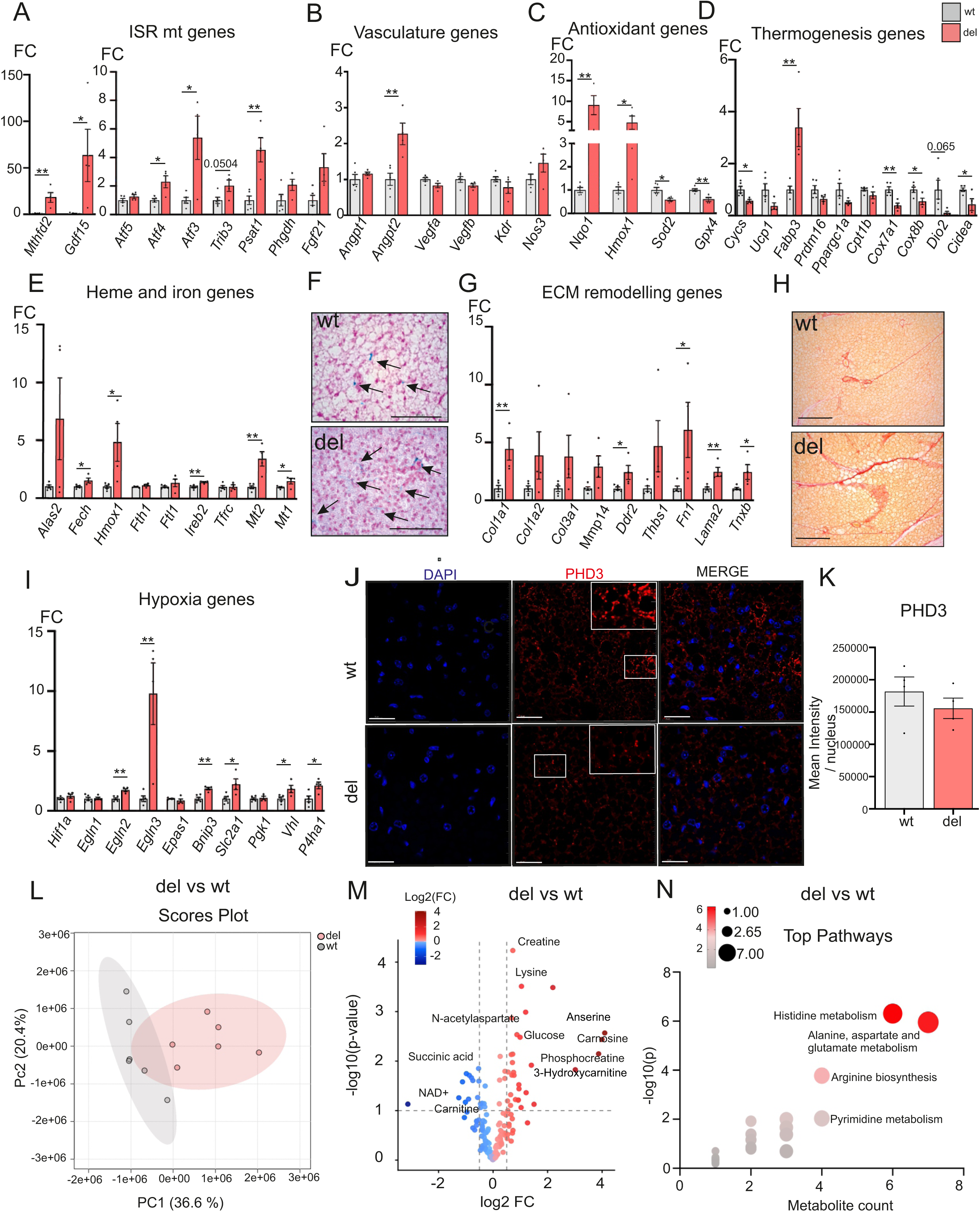
Transcriptomic analysis reveals coordinated pseudohypoxic, iron, stress, vascular, and extracellular matrix rewiring in Deletor BAT. (A) Transcript expression of mitochondrial integrated stress response (ISRmt) genes in WT and Deletor BAT, including *Mthfd2, Gdf15, Atf5, Atf4, Atf3, Trib3, Psat1, Phgdh,* and *Fgf21*. (n=5 (WT) and n = 4 (Del)) (B) Transcript expression of vascular modelling genes in WT and Deletor BAT, including *Angpt1, Angpt2, Notch1, Pecam1, Vegfa, Vegfb, Podxl, Flt1, Kdr, and Nos3.* (n=5 (WT) and n = 4 (Del)) (C) Transcript expression of antioxidant genes in WT and Deletor BAT, including *Nqo1, Hmox1, Sod2* and *Gpx4.* (n=5 (WT) and n = 4 (Del)) (D) Transcript expression of thermogenesis-associated genes in WT and Deletor BAT, including *Cycs, Ucp1, Fabp3, Prdm16, Ppargc1a, Cpt1b, Cox7a1, Cox8b, Dio2,* and *Cidea.* (n=5 (WT and n = 4 (Del)) (E) Transcript expression of iron and heme metabolism genes in WT and Deletor BAT, including *Alas2, Fech, Hmox1, Fth1, Ftl1, Ireb2, Tfrc, Mt2, and Mt1.* (n=5 (WT) and n = 4 (Del)) (F) Perl’s Prussian Blue staining of BAT sections from WT and Deletor mice showing increased ferric iron accumulation in Deletor BAT. Scale bar = 60 µm. (n=4 (WT) and n = 4 (Del)) (G) Transcript expression of extracellular matrix remodelling genes in WT and Deletor BAT, including *Col1a1, Col1a2, Col3a1, Mmp14, Ddr2, Thbs1, Fn1, Lama2, and Tnxb. (*n=5 (WT) and n = 4 (Del)) (H) Representative Picrosirius Red staining of BAT sections from wild-type (WT) and Deletor (Del) mice showing increased collagen deposition in Deletor BAT. Scale bar = 200 µm (I) Gene expression of hypoxia-associated genes in wild-type (WT) and Deletor (Del) BAT, including *Hif1a, Egln1, Egln2, Egln3, Epas1, Bnip3, Slc2a1, Pgk1, Vhl*, and *P4ha1* (n=5 (WT) and n = 4 (Del)). (J) Representative immunofluorescence images of BAT sections stained for DAPI and PHD3 (EGLN3). Scale bar = 20 µm (K) Quantification of PHD3 staining intensity per nucleus in wild-type (WT) and Deletor (Del) BAT (n=4 per group). (L) Principal component analysis of targeted metabolomics data from wild-type (WT) and Deletor (Del) BAT, showing clear separation of the two groups. (M) Volcano plot of differentially abundant metabolites in Deletor versus wild type BAT, with selected metabolites highlighted. (n=6 per group). (N) Metabolite pathway enrichment analysis showing the top significantly altered pathways in Deletor BAT Data are shown as mean ± SEM with individual data points overlaid where applicable. Statistical significance was assessed by unpaired Student’s t-test. *p<0.05, **p<0.01. For image quantification, 5 images were acquired from each animal and average was used for the analysis. (RNA seq data is analysed from WT = 5 and Del = 4 mice). Data are shown as mean ± SEM with individual data points overlaid where applicable. Statistical significance was assessed by unpaired Student’s t-test. *p<0.05, **p<0.01. For image quantification, 5 images were acquired from each animal and average was used for the analysis.

Genes involved in iron and heme metabolism were coordinately upregulated in Deletor BAT, including *Alas2, Fech, Hmox1, Fth1, Ftl1, Ireb2, Tfrc, Mt1,* and *Mt2* (Fig. 2E), indicating remodelling across heme synthesis, iron acquisition, iron-regulatory, and iron-buffering pathways. Perl’s Prussian Blue staining confirmed increased ferric iron accumulation in Deletor BAT at the tissue level (Fig. 2F), corroborating the transcriptional iron stress signature and iron dysregulation is a structural as well as a molecular feature of the disease state. This iron-stress program was accompanied by a selective redox-response signature. Deletor BAT showed marked induction of the stress-responsive genes *Nqo1* and *Hmox1*, with *Nqo1* increased approximately 9-fold and *Hmox1* approximately 5-fold relative to wild-type controls (Fig. 2C). In contrast, *Sod2*, which encodes mitochondrial superoxide dismutase, and *Gpx4*, which protects against lipid peroxide accumulation, were significantly reduced in Deletor BAT (Fig. 2C). This pattern indicates activation of cytoprotective oxidative stress-response pathways together with reduced mitochondrial antioxidant and lipid-peroxide detoxification capacity. Together with ferric iron accumulation, these findings suggest that Deletor BAT is vulnerable to oxidative and iron-linked lipid stress, although ferroptotic injury was not directly assessed. Extracellular matrix remodelling program was also evident, with significant upregulation of *Col1a1, Ddr2, Fn1, Lama2,* and *Tnxb* and an increasing trend in *Col1a2, Col3a1* and *Thbs1* (Fig. 2G), accompanied by Picrosirius Red staining confirming increased collagen deposition in Deletor BAT (Fig. 2H), demonstrating that the ECM transcriptional program is accompanied by overt structural fibrotic remodelling of the tissue.

Interestingly, hypoxia-associated transcriptional program was also prominently induced in the absence of environmental oxygen deprivation. Egln3 showed the most striking induction among hypoxia-associated genes, with approximately 10-fold elevation relative to wild-type controls, accompanied by significantly increased expression of *Egln2, Bnip3, Slc2a1, Vhl, and P4ha1* (Fig. 2I), supporting activation of HIF-mediated signalling secondary to respiratory chain dysfunction. PHD3 immunostaining revealed no significant change in PHD3 protein abundance per nucleus in Deletor BAT relative to wild-type controls (Fig. 2J, K), suggesting functional impairment rather than a change in protein level, consistent with a model in which PHD3 cofactor insufficiency, rather than loss of protein expression, underlies this signalling. Together, these transcriptional and histological data establish that Deletor BAT activates a highly structured disease program encompassing ISRmt activation, pseudohypoxic signalling, iron and heme pathway dysregulation, thermogenesis suppression, vascular expansion, and extracellular matrix remodelling identifying a pseudohypoxia–iron stress state as a defining molecular feature of BAT pathology in mitochondrial disease.

### Mitochondrial dysfunction drives coordinated TCA, glycolytic, and amino acid metabolic rewiring in brown adipose tissue

To define the biochemical consequences of chronic mtDNA instability in brown adipose tissue, we performed targeted metabolomic profiling of Deletor and wild-type BAT, quantifying 168 metabolites. Principal component analysis demonstrated clear separation of the two genotypes along PC1 (36.6%), indicating that mitochondrial dysfunction imposes a distinct remodelling of the BAT metabolome (Fig. 2L). Differential metabolite analysis identified significantly altered metabolites in Deletor BAT relative to wild-type controls (Fig. 2M). Pathway enrichment analysis highlighted histidine metabolism, arginine biosynthesis, alanine–aspartate–glutamate metabolism, and pyrimidine metabolism among the most significantly altered pathways (Fig. 2N), consistent with broad disruption of amino acid handling, nucleotide metabolism, and central carbon metabolism downstream of respiratory chain dysfunction.

Deletor BAT displayed signature of altered nutrient handling and impaired mitochondrial substrate. Lysine, histidine, arginine, and asparagine were significantly elevated, while alanine showed an increasing trend, whereas aspartate and cysteine were reduced (Fig. 3A), consistent with impaired amino acid catabolism and altered routing of amino acid-derived carbon and nitrogen substrates under conditions of mitochondrial dysfunction. Creatine and phosphocreatine were also elevated in Deletor BAT (Fig. 3B), indicating remodelling of energy-buffering pathways in response to respiratory chain impairment, consistent with reduced downstream ATP utilisation capacity. In parallel, glucose was significantly increased, with ribose-5-phosphate and fructose-6-phosphate showing an increasing trend, while glycerol-3-phosphate was reduced (Fig. 3C), consistent with accumulation of glycolytic and pentose phosphate pathway intermediates alongside impaired glycerol-3-phosphate shuttle activity and reduced entry of glucose-derived carbon into mitochondrial oxidation.

**Figure 3.**
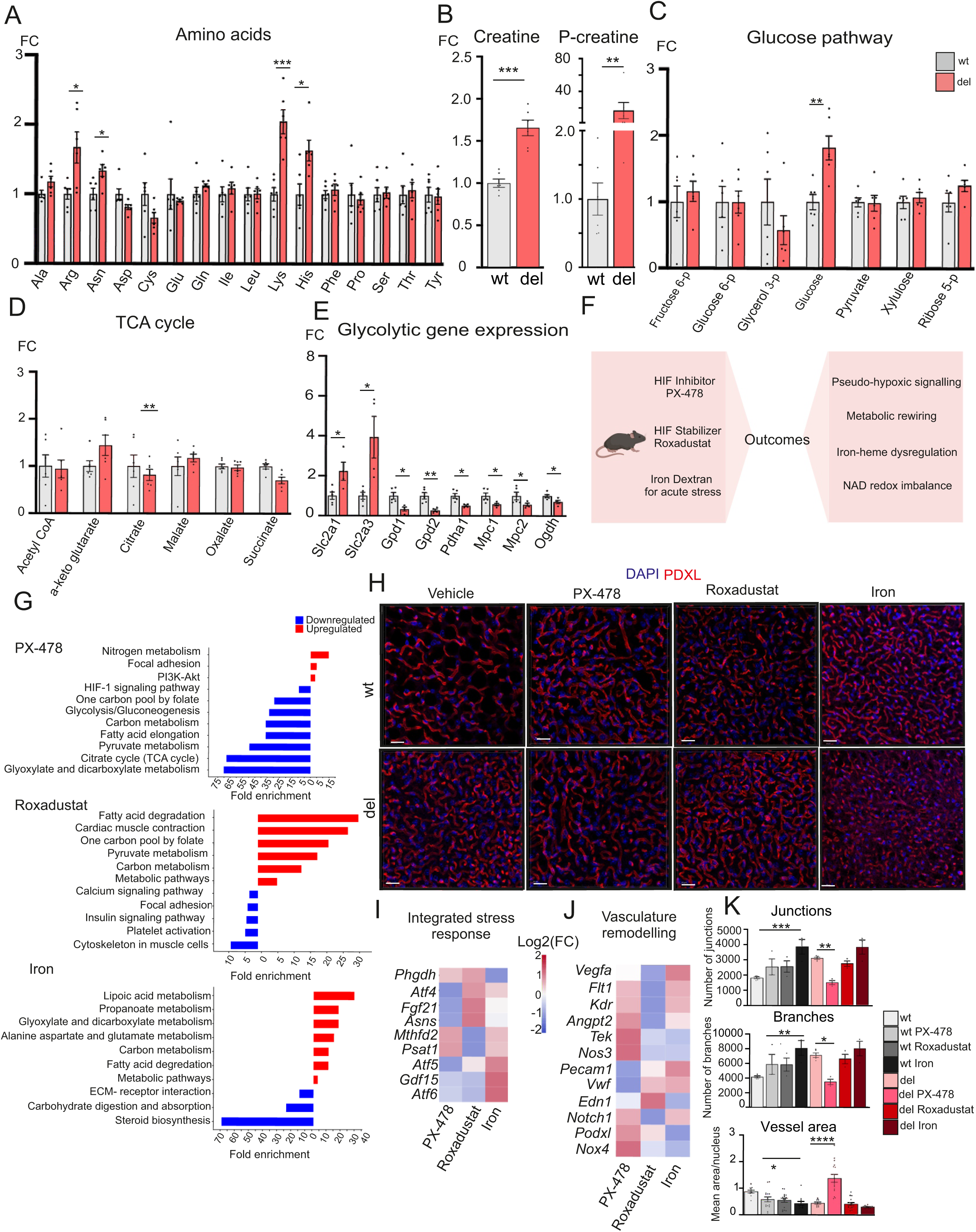
Metabolic rewiring in BAT. (A) Relative abundance of selected amino acids in wild-type (WT) and Deletor (Del) BAT, showing accumulation of several amino acids including arginine, lysine, histidine, and asparagine, together with reduction of aspartate and cysteine (n=6 per group). (B) Relative abundance of creatine and phosphocreatine in wild-type (WT) and Deletor (Del) BAT, indicating altered energy-buffering metabolism in Deletor BAT. (n=6 per group) (C) Relative abundance of glucose-associated metabolites, including fructose-6-phosphate, glucose-6-phosphate, glycerol-3-phosphate, glucose, pyruvate, xylulose, and ribose-5-phosphate, showing accumulation of glycolytic and glucose-pathway intermediates in Deletor BAT (n=6 per group). (D) Relative abundance of selected TCA cycle intermediates in wild-type (WT) and Deletor (Del) BAT, including acetyl-CoA, α-ketoglutarate, citrate/isocitrate, malate, oxalate, and succinate, showing non-uniform remodelling (n=6 per group). (E) Expression of genes involved in glucose uptake, glycerol phosphate metabolism, pyruvate handling, and mitochondrial carbon oxidation, including *Slc2a1, Slc2a3, Gpd1, Gpd2, Pdha1, Mpc1, Mpc2 and Ogdh* (n=5 (WT) and n = 4 (Del)). (F) Experimental design for pharmacological perturbation of Deletor BAT. Wild-type and Deletor mice were treated with PX-478, roxadustat, or iron dextran, and BAT transcriptomes were compared with vehicle-treated Deletor controls. (G) KEGG pathway enrichment analysis of differentially expressed genes in Deletor BAT following PX-478, roxadustat, or iron dextran treatment relative to Deletor vehicle controls. Upregulated and downregulated pathways are shown for each treatment. (H) Representative podocalyxin immunofluorescence staining of BAT sections from vehicle-, PX-478-, roxadustat-, and iron dextran-treated Deletor (Del) and wild-type (WT) mice, showing treatment-specific changes in vascular architecture. Scale bar = 20 µm. (I) Heatmap showing treatment-dependent regulation of mitochondrial integrated stress response genes, including *Phgdh, Atf4, Fgf21, Asns, Mthfd2, Psat1, Atf5, Gdf15*, and *Atf6* across PX-478, roxadustat, and iron dextran treatments (n=5 per group). (J) Heatmap showing treatment-dependent regulation of vascular remodelling genes, including *Vegfa, Flt1, Kdr, Angpt2, Tek, Nos3, Pecam1, Vwf, Edn1, Notch1, Podxl,* and *Nox4* (n=5 per group). (K) Quantification of vascular branches, junctions (n=5 (WT and Del), others n=3 per group), and vessel area in BAT sections across treatment groups. Data are shown as mean ± SEM with individual data points overlaid where applicable. Statistical Student’s t-test assessed statistical significance. For vascular analysis, statistical significance was assessed by one-way ANOVA followed by Tukey’s multiple comparisons test. *p<0.05, **p<0.01, ***p<0.001, ****p<0.0001. For image quantification, 5 images were acquired from each animal, and average was used for the analysis.

TCA cycle intermediates were altered in a non-uniform pattern, Deletor BAT showed increased α-ketoglutarate and malate, alongside reduced citrate and succinate (Fig. 3D). This pattern is consistent with selective disruption of mitochondrial carbon flux rather than global depletion of TCA cycle metabolites and indicates impaired oxidative metabolism and altered redox-dependent dehydrogenase activity downstream of respiratory chain dysfunction. Consistent with this metabolic shift, transcriptomic analysis revealed coordinated remodelling of genes governing glucose handling and mitochondrial carbon entry in Deletor BAT (Fig. 3E). In contrast, *Gpd1*, *Gpd2*, *Pdha1*, *Mpc1*, *Mpc2* and *Ogdh* were suppressed, reflecting impaired mitochondrial pyruvate entry, reduced TCA cycle activity, and decreased oxidative carbon utilisation (Fig. 3E). This pattern indicates that glucose uptake capacity is increased while mitochondrial oxidation capacity is transcriptionally constrained. Consistent with this, indirect calorimetry during acute cold challenge revealed impaired fuel switching in Deletor mice: whereas wild-type mice reduced RER toward lipid oxidation, Deletor mice maintained an elevated RER near 1.0, indicating defective transition toward lipid-based fuel utilisation under thermogenic demand (Supp Fig. 1D). Together, these findings demonstrate that chronic mtDNA instability shifts BAT away from the high oxidative capacity required for thermogenic function and toward an energy-stressed, glycolytically remodelled metabolic state, accompanied by impaired cold-adaptive substrate switching.

### Pharmacological HIF-axis modulation reveals maladaptive stress and vascular remodelling in Deletor BAT

To determine how modulation of pseudohypoxic signalling and iron availability reshapes BAT pathology, wild-type and Deletor mice were treated with the HIF-1α inhibitor PX-478, the HIF-stabilising pan-PHD inhibitor roxadustat, or iron dextran to introduce iron stress. PX-478 and roxadustat were used to bidirectionally modulate the PHD3–HIF axis, while iron dextran was administered to determine whether iron loading alone is sufficient to recapitulate key features of the Deletor BAT disease remodelling. BAT transcriptomes from each treatment arm were compared with vehicle-treated Deletor controls (Fig. 3F). KEGG pathway enrichment analysis revealed distinct treatment-dependent transcriptional responses across the three interventions (Fig. 3G). PX-478 suppressed glycolysis, TCA cycle, and one-carbon metabolism pathways, consistent with attenuation of the pseudohypoxic metabolic stress program. Roxadustat upregulated fatty acid degradation and carbon metabolism while downregulating adrenergic and calcium signalling, consistent with HIF stabilisation-driven remodelling. Iron dextran induced carbon metabolism, fatty acid metabolism, and fatty acid elongation pathways alongside suppression of ECM-receptor interaction programs, consistent with iron-driven metabolic and structural stress.

Interestingly, PX-478 markedly attenuated the ISRmt program in Deletor BAT, with reduced expression of *Gdf15, Fgf21, Atf4, Atf5*, and *Asns* (Fig. 3I), demonstrating that HIF-1α suppression attenuates the maladaptive stress-response program. Both roxadustat and iron produced a limited effect or increasing trend on ISRmt genes. In contrast, vascular remodelling genes including *Flt1, Kdr, Angpt2, Tek, Nos3, Notch1, Podxl,* and *Nox4* were induced by PX-478 at the transcriptional level (Fig. 3J). Podocalyxin immunostaining revealed that PX-478 treatment reduced vascular branching and junction complexity but increased mean vessel area in Deletor BAT (Fig. 3H), indicating that HIF-1α suppression remodels vascular architecture toward fewer but larger vessels. Roxadustat and iron dextran had no significant effect on vascular branch count, junction density, or vessel area relative to vehicle-treated Deletor BAT. Representative three-dimensional Imaris reconstructions further illustrated alterations in vascular network architecture across the experimental groups. (Supp. Fig. 2B). Quantification confirmed these treatment-specific vascular effects (Fig. 3K), demonstrating that the hyperbranched vascular network of Deletor BAT is at least partly HIF-dependent and that HIF suppression partially normalises vascular architecture despite inducing a compensatory transcriptional angiogenic response.

Adipocyte morphology was not significantly altered by any intervention (Supp. Fig. 2A). UCP1 immunoreactivity remained reduced in PX-478 and roxadustat-treated Deletor BAT and was further suppressed by iron (Supp. Fig. 3A), demonstrating that thermogenic identity cannot be restored by targeting the pseudohypoxic program alone and that iron loading independently compounds to thermogenic failure.Supp

Together, these data demonstrate that ISRmt activation and vascular remodelling are pharmacologically separable components of Deletor BAT pathology. HIF-1α inhibition attenuates the stress-response program and partially normalises vascular architecture of Deletor BAT reducing hyperbranching and junction complexity while restoring vessel area toward wild-type dimensions. Roxadustat and iron dextran each reinforce disease-associated features, with iron additionally driving thermogenic suppression independently of the pseudohypoxia program, collectively establishing that pseudohypoxia signalling is causally involved in both the stress and vascular components of Deletor BAT pathology.

### HIF-axis modulation and iron loading reveal pharmacologically separable disease programs in Deletor BAT

To determine how pharmacological modulation of HIF signalling and iron availability affects tissue remodelling in Deletor BAT, we examined extracellular matrix deposition, iron metabolism, pseudohypoxic signalling, glucose metabolism and immune infiltration across treatment arms.

Extracellular matrix remodelling genes remained elevated across all Deletor treatment groups (Fig. 4A). Neither PX-478 nor roxadustat attenuated this program, whereas iron dextran produced the strongest fibrotic transcriptional response. Picrosirius Red staining confirmed these findings at the tissue level (Fig. 4B): collagen accumulation persisted after PX-478 and roxadustat treatment, while iron dextran induced prominent perivascular and interstitial collagen deposition in both control and Deletor mice. Iron/heme genes showed more treatment-specific regulation (Fig. 4C). PX-478 attenuated this stress program, roxadustat maintained or enhanced it, and iron dextran produced the strongest induction of storage and metal-stress genes, including increased expression of *Fth1, Mt1, Mt2,* and *Lcn2*. Among these, FTH1, the ferritin heavy chain and principal iron-storage protein - serves as a direct marker of cellular iron buffering capacity, converting reactive labile iron into a safely sequestered form. FTH1 protein followed the same overall pattern (Fig. 4D): low in wild-type BAT, elevated in vehicle-treated Deletor BAT, reduced toward wild-type levels by PX-478, further increased by roxadustat, and maximally induced by iron dextran. Perls Prussian Blue staining corroborated this pattern at the tissue level, showing treatment-dependent iron accumulation with iron dextran producing the strongest ferric iron deposition in control and Deletor mice (Supp. Fig. 3B), consistent with iron dextran driving the strongest iron-storage response and the most severe fibrotic phenotype.

**Figure 4.**
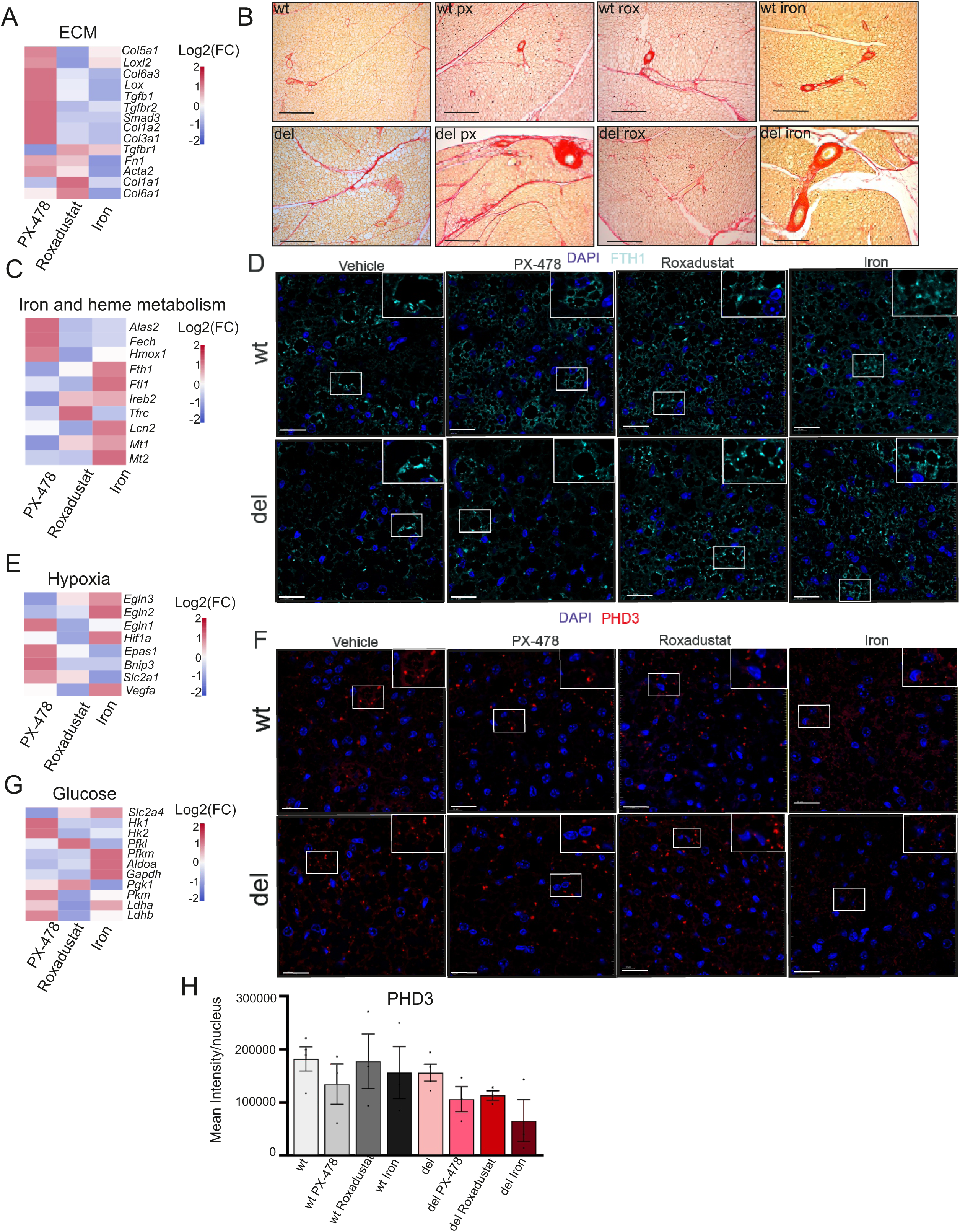
HIF-axis modulation and iron loading produce divergent ECM, iron, pseudohypoxic, and glycolytic remodelling programs in Deletor BAT. (A) Heatmap showing expression of extracellular matrix and fibrosis-associated genes across wild-type and Deletor BAT treatment groups, including *Col5a1, Loxl2, Col6a3, Lox, Tgfb1, Tgfbr2, Smad3, Col1a2, Col3a1, Tgfbr1, Fn1, Acta2, Col1a1,* and *Col6a1*. (n=5 per group) (B) Representative Picrosirius Red staining of BAT sections from vehicle-, PX-478-, roxadustat-, and iron dextran-treated groups, showing collagen deposition and fibrotic remodelling in Deletor and WT BAT. Scale bar = 200 µm (C) Heatmap showing expression of iron and heme metabolism genes, including *Alas2, Fech, Hmox1, Fth1, Ftl1, Ireb2, Tfrc, Lcn2, Mt1,* and *Mt2*, across treatment groups. (n=5 per group) (D) Representative immunofluorescence images of BAT sections stained for DAPI and FTH1, showing treatment-dependent regulation of ferritin heavy chain protein abundance. Scale bar = 20 µm. (E) Heatmap showing expression of HIF/pseudohypoxia-associated genes, including *Egln3, Egln2, Egln1, Hif1a, Epas1, Bnip3, Slc2a1*, and *Vegfa,* across treatment groups. (n=5 per group) (F) Representative immunofluorescence images of BAT sections stained for DAPI and PHD3, in Deletor BAT and differential regulation following PX-478, roxadustat, and iron dextran treatment. Scale bar = 20 µm (G) Heatmap showing expression of glucose transport and glycolytic genes, including *Slc2a4, Hk1, Hk2, Pfkl, Pfkm, Aldoa, Gapdh, Pgk1, Pkm, Ldha,* and *Ldhb,* across treatment groups. (n=5 per group) (H) Quantification of PHD3 staining intensity per nucleus in BAT sections across treatment groups (n=4 (WT and Del), others n=3 per group). Heatmaps show normalised gene expression values. Representative images are shown for each treatment group. Data are shown as mean ± SEM with individual data points overlaid where applicable. Statistical significance was assessed by one-way ANOVA followed by Tukey’s multiple comparisons test. For image quantification, 5 images were acquired from each animal and average was used for the analysis.

The pseudohypoxia-associated program was also treatment-responsive (Fig. 4E). PX-478 reduced expression of HIF-responsive genes, including *Egln3*, whereas roxadustat maintained or increased this program. Iron dextran produced a distinct response, indicating that iron loading can sustain components of the disease state without fully phenocopying HIF stabilisation. At the protein level, PHD3 immunostaining showed treatment-specific changes relative to vehicle-treated Deletor BAT (Fig. 4F, H): PX-478 and roxadustat each produced a mild reduction in PHD3 protein, whereas iron dextran produced a more prominent reduction. Glycolytic genes followed a similar treatment-dependent pattern (Fig. 4G), with PX-478 attenuating and roxadustat sustaining glycolytic remodelling.

Because iron loading and fibrotic remodelling are frequently accompanied by immune cell recruitment, we next asked whether innate immune infiltration followed a similar treatment-dependent pattern in Deletor BAT. CD68 staining showed sparse, mainly perivascular macrophages in wild-type BAT, whereas vehicle-treated Deletor BAT showed increased parenchymal CD68+ cells (Supp. Fig. 4A). PX-478 reduced CD68+ cell density, roxadustat and iron-maintained vehicle-level accumulation. KEGG analysis in wild-type mice exposed to the same treatments did not show enrichment of the disease-defining programs, including ISRmt, pseudohypoxia, ECM remodelling, or immune recruitment (Supp. Fig. 4B), indicating that these responses are largely disease-context dependent rather than generic drug effects. Together, these data show that the disease-associated remodelling modules in Deletor BAT are pharmacologically separable. PX-478 suppresses pseudohypoxic, iron glycolytic, and immune stress programs but does not resolve established ECM remodelling. Roxadustat reinforces HIF and iron-linked disease features, whereas iron dextran drives the strongest iron-storage, inflammatory, and fibrotic response. Thus, pseudohypoxic signalling, iron stress, glycolytic remodelling, and immune infiltration remain treatment-responsive, whereas collagen deposition is comparatively refractory once established.

### Nicotinamide riboside restores NAD^+^ redox balance and broadly attenuates metabolic and transcriptional stress in Deletor BAT

Deletor BAT showed a striking imbalance in NAD⁺ pathway gene expression: core biosynthesis and salvage genes, including *Nampt, Nadk, Sirt3,* and *Slc25a51*, were suppressed, while NAD⁺-consuming and compensatory enzymes, including *Nmrk2, Nnmt, Parp8, Parp3,* and *Cd38*, were significantly induced (Fig. 5A). *Nmrk2* and Parp8 showed the most significant induction in the entire NAD gene set, with *Nmrk2* elevated more than 30-fold above wild-type levels, indicating that Deletor BAT mounts a profound compensatory response to NAD⁺ depletion while simultaneously losing the biosynthetic capacity to restore it. This pattern positions pyridine nucleotide homeostasis as a critically stressed node embedded within the pseudohypoxic remodelling program of Deletor BAT and motivated direct therapeutic interrogation of the NAD⁺ axis. Deletor and wild-type mice were therefore treated for six months with nicotinamide riboside (NR; 400 mg/kg/day), a biosynthetic NAD⁺ precursor, and BAT was subjected to integrated transcriptomic and metabolomic analysis (Fig. 5B). Deletor BAT exhibited a pronounced imbalance in pyridine nucleotide pools, characterised by significantly reduced NAD⁺, accumulation of NADH, and a markedly compressed NAD⁺/NADH ratio relative to wild-type controls (Fig. 5C). NR treatment robustly normalised all three parameters: increasing NAD⁺, reducing NADH accumulation, and restoring the NAD⁺/NADH ratio toward control values, demonstrating that the redox imbalance in Deletor BAT is fully reversible by exogenous NAD⁺ precursor supplementation (Fig. 5C).

**Figure 5.**
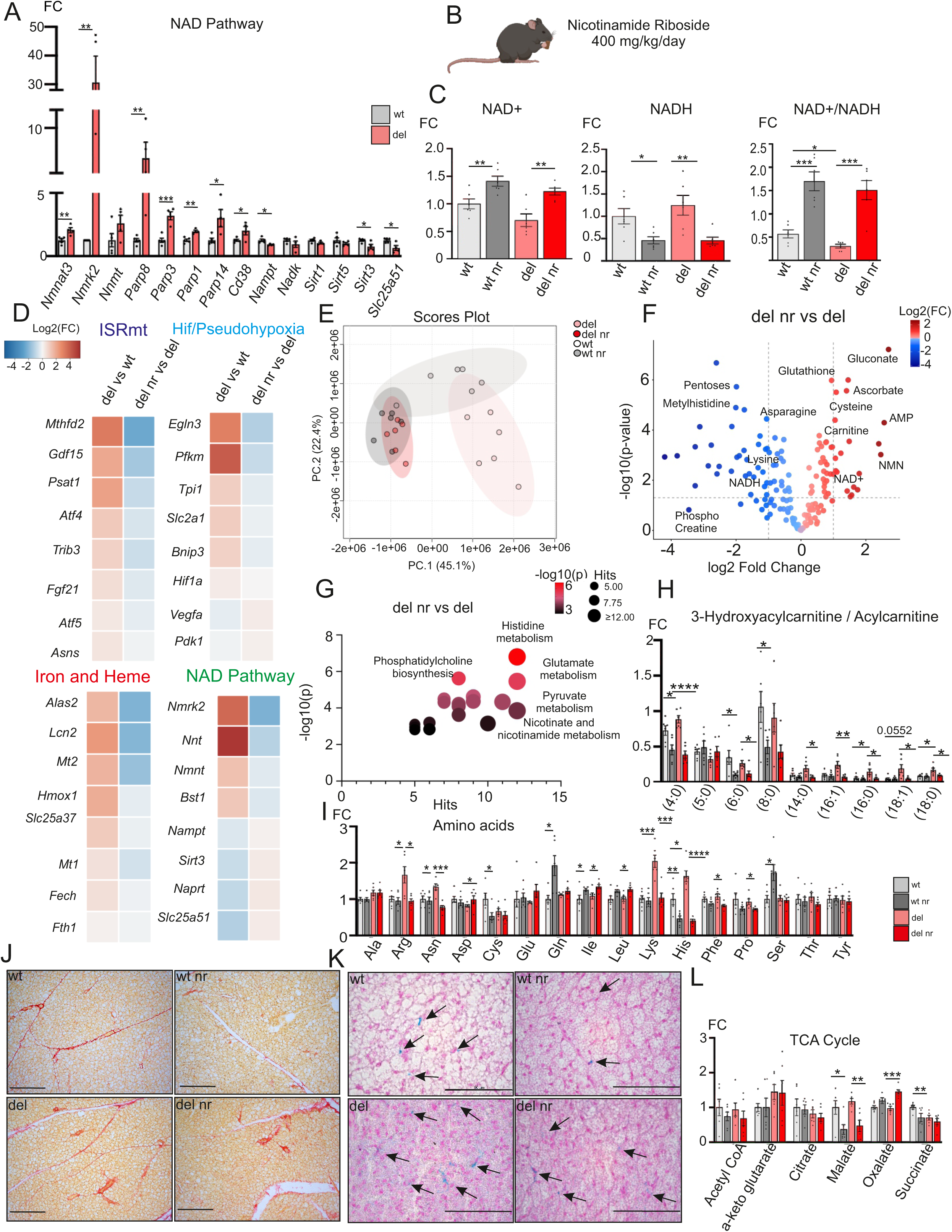
Nicotinamide riboside restores NAD+ redox balance and attenuates disease features. (A) Expression of NAD+ biosynthesis, salvage, utilisation, and transport-associated genes in wild-type (WT) and Deletor (Del) BAT, including *Nmnat3, Nmrk2, Nnmt, Parp8, Parp3, Parp1, Parp14, Cd38, Nampt, Nadk, Sirt1, Sirt5, Sirt3*, and *Slc25a51*. (n=5 (WT) and n= 4 (Del)) (B) Schematic of the nicotinamide riboside (NR) intervention. Wild type (WT) and Deletor (Del) mice were treated with NR for six months at 400 mg/kg/day, followed by BAT metabolomic and transcriptomic analysis. (C) Quantification of NAD+, NADH, and the NAD+/NADH ratio in BAT from WT vehicle, WT NR, Deletor vehicle, and Deletor NR mice, showing restoration of pyridine nucleotide redox balance following NR treatment (n= 6 per group). (D) Heatmap comparing disease-induced transcriptional changes and NR-responsive changes across four pathway modules: mitochondrial integrated stress response (ISRmt), iron and heme metabolism, HIF/pseudohypoxia, and NAD pathway genes. Values show relative transcriptional changes in Deletor vehicle versus WT vehicle and Deletor NR versus Deletor vehicle comparisons. (E) Principal component analysis of targeted metabolomics data from WT vehicle, WT-NR, Deletor vehicle, and Deletor-NR BAT, showing NR-dependent remodelling of the Deletor BAT metabolome (n=6 per group). (F) Volcano plot showing differentially abundant metabolites in Deletor NR BAT relative to Deletor vehicle BAT. Metabolites increased after NR treatment are shown in red and metabolites reduced after NR treatment are shown in blue (n=6 per group). (G) Metabolite pathway enrichment analysis of NR-responsive metabolites in Deletor BAT, highlighting histidine metabolism, glutamate metabolism, pyruvate metabolism, nicotinate and nicotinamide metabolism, and phosphatidylcholine biosynthesis. Bubble size indicates the number of pathway hits and colour intensity indicates statistical significance (n=6 per group). (H) Ratio of 3-hydroxyacylcarnitine to acylcarnitine across measured chain lengths (C4:0, C5:0, C6:0, C8:0, C14:0, C16:1, C16:0, C18:1, C18:0) in BAT from WT vehicle, WT NR, Deletor vehicle, and Deletor NR mice (n=6 per group). (I) Relative abundance of selected amino acids in WT vehicle, WT-NR, Deletor vehicle, and Deletor-NR BAT, showing partial normalisation of the amino acid stress signature following NR treatment (n=6 per group). (J) Representative Picrosirius Red staining of BAT sections from wt vehicle, wt nr, Deletor vehicle, and Deletor nr mice. Scale bar = 200 µm (K) Representative Perl’s Prussian Blue staining of BAT sections from wt vehicle, wt nr, Deletor vehicle, and Deletor nr mice. Scale bar = 60 µm. (L) Relative abundance of selected TCA cycle intermediates, including acetyl-CoA, α-ketoglutarate, citrate, malate, oxalate, and succinate, in WT vehicle, WT-NR, Deletor vehicle, and Deletor-NR BAT (n=6 per group). Data are shown as mean ± SEM with individual data points overlaid where applicable. Statistical significance was assessed by unpaired Student’s t-test (A) or one-way ANOVA followed by Tukey’s multiple comparisons test (C–J), as indicated. *p<0.05, **p<0.01, ***p<0.001, ****p<0.0001.

To define the transcriptional basis of NR-mediated rescue, RNA sequencing was performed in Deletor NR versus Deletor vehicle BAT. Pathway-level analysis using a four-module heatmap covering ISRmt, HIF/pseudohypoxia, iron and heme, and NAD pathway genes revealed coordinated attenuation of the disease transcriptional signature across all four modules following NR treatment (Fig. 5D). The ISRmt program showed the strongest rescue, with consistent downregulation of *Mthfd2, Gdf15, Atf4, Trib3, Psat1, Fgf21, Atf5*, and *Asns*, indicating broad suppression of the ATF4/ATF5 linked stress axis. Iron and heme stress genes, including *Alas2, Lcn2, Mt2, and Mt1*, were similarly attenuated, consistent with partial normalisation of iron stress downstream of NAD⁺ repletion. The HIF/pseudohypoxia program showed partial rescue: *Egln3*/PHD3 and *Pfkm* trended downward, indicating attenuation but not complete resolution of the pseudohypoxic program. The NAD+ pathway block revealed a bidirectional rescue pattern: disease-induced compensatory genes *Nmrk2, Nnt, Nnmt,* and *Bst1* were suppressed by NR, consistent with negative feedback from a restored NAD⁺ pool, while disease-depleted genes *Nampt, Sirt3, Naprt*, and *Slc25a51* trended toward recovery, indicating partial restoration of the core NAD⁺ biosynthetic and transport machinery.

Principal component analysis of the targeted metabolome demonstrated that NR treatment broadly restored the Deletor metabolic profile. Disease NR samples separated from Disease vehicle along PC1 (45.1%) and clustered with Control NR mice (Fig. 5E), indicating that NR reverses the disease-associated metabolic state at a global level rather than correcting isolated metabolite abnormalities. Unsupervised hierarchical clustering of differentially regulated metabolites corroborated this pattern (Supp. Fig. 5A), with Disease NR samples clustering closely with Control NR and separating clearly from Disease vehicle. Metabolites elevated in disease, including creatine and nucleotide-related species, were normalised by NR, while depleted metabolites such as carnitine and NAD⁺ were restored toward control levels. Consistent with this, volcano plot analysis identified widespread metabolite changes in Deletor NR relative to Deletor vehicle BAT, with the majority of significantly altered metabolites showing reduced abundance following NR treatment (Fig. 5F), indicating broad suppression of the disease-associated stress metabolome. Pathway enrichment analysis identified histidine metabolism, phosphatidylcholine biosynthesis, glutamate metabolism, pyruvate metabolism, and nicotinate and nicotinamide metabolism as the most significantly rescued programs (Fig. 5G), with enrichment of nicotinate and nicotinamide metabolism providing independent metabolomic confirmation of successful NAD⁺ repletion in Deletor BAT. NR also normalised the disease-associated amino acid signature of Deletor BAT: lysine, arginine, asparagine, and histidine, each significantly elevated in Deletor BAT relative to wild-type controls, were normalised following treatment, while aspartate and cysteine, which were reduced in Deletor BAT, showed partial restoration (Fig. 5I). This is consistent with NR-mediated restoration of amino acid catabolism and mitochondrial substrate, rather than a non-specific global effect on amino acid metabolism. NR induced broad metabolic remodelling in both wild-type and Deletor BAT, indicating that NAD⁺ repletion has substantial metabolic effects in adipose tissue regardless of disease status with the Deletor-specific rescue reflecting correction of disease-associated metabolite abnormalities on top of these shared NR-induced changes (Supp Fig. 5 B, C)

Strikingly, the ratio of 3-hydroxyacylcarnitine to acylcarnitine a marker of limited activity at the NAD⁺-dependent 3-hydroxyacyl-CoA dehydrogenase step of β-oxidation was significantly elevated in Deletor BAT relative to wild-type at multiple chain lengths and was normalised toward wild-type levels following NR treatment (Fig. 5H). Absolute pool sizes of free carnitine and acylcarnitine species followed a broadly consistent pattern, with reduced levels in Deletor BAT restored toward wild-type following NR treatment (Supp. Fig. 5F). This rescue is mechanistically coherent with the coordinate normalisation by NR of lysine, α-ketoglutarate, and ascorbate, the substrates and cofactors of the carnitine biosynthesis pathway (Fig. 5I,L). Together, these findings link the resolution of this NAD⁺-dependent bottleneck to recovery of fatty acid handling capacity in Deletor BAT.

Despite this broad rescue, established structural remodelling remained largely refractory to NR treatment. Picrosirius Red staining confirmed persistent perivascular and interstitial fibrotic remodelling in NR-treated Deletor mice, comparable to vehicle-treated controls (Fig. 5J), and ECM-associated genes, including *Col1a1, Col1a2, Col3a1, Mmp14, Ddr2, Thbs1, Fn1, Lama2*, and *Tnxb,* remained increased (Supp. Fig. 5D). Similarly, thermogenesis-associated genes, including *Cycs, Ucp1, Fabp3, Prdm16, Ppargc1a, Cpt1b, Cox7a1, Cox8b, Dio2*, and *Cidea,* remained suppressed following NR treatment (Supp. Fig. 5E), indicating that loss of thermogenic identity is not downstream of NAD⁺ redox imbalance and cannot be restored by NAD⁺ repletion alone at this stage of disease. In contrast, Perls’ Prussian Blue staining revealed a reduction in ferric iron accumulation in Deletor NR BAT relative to Deletor vehicle controls (Fig. 5K), indicating that NR treatment partially normalises tissue iron burden alongside its transcriptional suppression of iron stress genes, connecting the metabolic rescue to a measurable reduction in iron-associated pathology at the histological level.

Together, these findings identify NAD⁺ redox imbalance as a central and metabolically reversible component of BAT dysfunction in Deletor mice. NR supplementation suppressed ISRmt, pseudohypoxic, and iron stress programs while restoring NAD⁺ redox balance, amino acid homeostasis, and carnitine-dependent fatty acid handling, and partially normalised tissue iron burden. However, established fibrotic remodelling and loss of thermogenic identity remained refractory to NR treatment, defining the structural limit of NAD⁺ supplementation in advanced mitochondrial BAT disease and motivating combination strategies targeting both the NAD⁺ axis and these structural endpoints.

## Discussion

Mitochondrial DNA maintenance disorders cause progressive multisystem degeneration ([8,30–33] driven by respiratory chain failure, but the contribution of brown adipose tissue, a tissue whose entire functional identity depends on intact mitochondrial respiration [34], to this disease spectrum has never been examined. Our study identifies BAT as a vulnerable oxidative tissue whose structure and function are fundamentally compromised when mitochondrial genome maintenance fails. Critically, this molecular and structural compromise translates to functional thermogenic failure under physiological demand, Whole-body indirect calorimetry showed reduced oxygen consumption during acute cold challenge and impaired RER-based fuel switching, consistent with impaired cold-adaptive oxidative metabolism and corroborating the NAD⁺-dependent β-oxidation bottleneck identified metabolically. Consistent with findings previously established in skeletal muscle, Deletor BAT activated a robust mitochondrial integrated stress response alongside coordinate upregulation of one-carbon pool by folate and serine biosynthesis pathways [11–13,35]. This shared stress architecture indicates that the ISRmt is a conserved tissue-level response to mtDNA deletion disease across metabolically distinct tissues.

A central finding is that Deletor BAT activates a pseudohypoxic stress program despite the absence of imposed hypoxia. Induction of *Egln3, Hif1a, Epas1*, and HIF-responsive metabolic genes supports a model in which respiratory chain dysfunction perturbs the intracellular determinants of oxygen sensing, including redox state, TCA metabolite availability, and iron handling rather than depleting environmental oxygen [17,19,20]. This is particularly important in BAT, where mitochondrial oxidation is not only a source of ATP but a defining feature of thermogenic identity [5]. Notably, the striking transcriptional induction of *Egln3 (Phd3)* was not matched at the protein level, where EGLN3 abundance was unchanged or only mildly reduced relative to control mice. EGLN3 itself is a HIF target gene, creating a feed-forward amplification loop once pseudohypoxic signalling is initiated [36,37], confirming activation of HIF1a. EGLN3 also has non-canonical, HIF-independent functions including suppression of fatty acid oxidation through ACC2 hydroxylation (Yoon et al, 2020; German et al, 2016), which may further compound thermogenic failure through mechanisms distinct from HIF activation itself. Further, EGLN3 catalysis requires ferrous iron as a cofactor [40], the coordinate shift toward ferric iron accumulation and Fth1 induction in Deletor BAT offers a plausible biochemical basis for this functional impairment[41]: expansion of the iron-storage pool at the expense of catalytically available Fe²⁺ would limit PHD3 hydroxylase activity without requiring any change in enzyme abundance. This reframes pseudohypoxia in Deletor BAT because of cofactor-level enzyme dysfunction rather than a purely transcriptional phenomenon, positioning PHD3 as a molecular integrator of respiratory failure and iron mishandling into a self-reinforcing pseudohypoxic state.

The accumulation of creatine and phosphocreatine in Deletor BAT may reflect a compensatory attempt to engage UCP1-independent creatine futile cycling, an alternative thermogenic mechanism recently described in BAT [42–44]. The NR-responsive correction of carnitine and acylcarnitine abnormalities is consistent with improved NAD⁺-dependent fatty-acid handling. Together, these findings further support the view that Deletor BAT mounts extensive metabolic adaptations that ultimately fail to preserve thermogenic identity. Pharmacological bidirectional modulation of the PHD3-HIF axis provided functional evidence that this pseudohypoxic program is maladaptive rather than compensatory. Using the HIF-1α inhibitor PX-478 [45,46] and the clinically approved PHD inhibitor roxadustat [47,48]as mechanistic opposites revealed a striking asymmetry: HIF-1α suppression attenuated the mitochondrial integrated stress response, suppressed iron and pseudohypoxic gene programs, and partially normalised vascular architecture, whereas roxadustat produced no benefit and in several respects worsened disease features. This asymmetry constitutes causal evidence that PHD3-HIF signalling actively drives the maladaptive stress state in Deletor BAT, rather than passively reflecting tissue damage. Unlike tissues in which HIF-driven glycolysis supports survival under genuine oxygen limitation [49–51], BAT cannot tolerate a chronic shift away from mitochondrial oxidation without compromising its core physiological function; HIF activation in this context does not rescue the tissue but accelerates its functional decompensation. Systemic iron loading with iron dextran provided a third and mechanistically independent line of evidence recapitulating and in several respects exacerbating disease features including the strongest fibrotic response and further suppression of thermogenic identity. In a tissue whose respiratory machinery depends entirely on iron-sulphur clusters, haem synthesis, and electron transport chain assembly ([52–54] redistribution of iron into stress-associated pools simultaneously impairs oxidative capacity and promotes fibrotic remodelling. The recent identification of a haem and branched chain amino acids driving BAT thermogenesis further underscores the unique vulnerability of brown adipose tissue to iron and haem pathway disruption [23].

NAD⁺ redox imbalance emerged as the most therapeutically tractable node in the Deletor BAT disease network, and nicotinamide riboside as the most broadly effective intervention tested. This suggests that Deletor BAT phenotype depends on downstream redox crisis that remains correctable even when the upstream mtDNA deletion did not change, consistent with prior work identifying NAD⁺ depletion as a common and therapeutically tractable node across mitochondrial disease mice models [28,55,56] and myopathy patients ([28,29]. The chain-length-selective normalisation of the 3-hydroxyacylcarnitine to acylcarnitine ratio following NR treatment provides biochemical support for this model, indicating that NAD⁺ repletion acts, at least in part, through a specific and identifiable enzymatic node rather than a more diffuse metabolic effect [57,58]. NR was able to rescue key transcriptional changes, spanning ISRmt, pseudohypoxic, and iron-stress programs simultaneously suggesting that these three programs are not independent parallel injuries but are mechanistically coupled downstream of a shared NAD⁺ dependent redox state. However, NR did not restore thermogenic identity or reverse established fibrosis, indicating that some downstream endpoints become treatment-refractory once structural remodelling is consolidated. These findings position NAD^+^ repletion as a strategy most likely to be effective early in disease, before irreversible loss of cellular identity and fibrotic remodelling limit tissue recovery.

In summary, this study identifies brown adipose tissue as a previously underappreciated target of mitochondrial genome maintenance disease and defines a pseudohypoxia-iron-NAD^+^ axis as a central organiser of its pathology. The persistence or worsening of disease features after roxadustat highlights a potential safety consideration for PHD inhibition in mitochondrial dysfunction; the partial rescue achieved by PX-478 supports functional involvement of the pseudohypoxic program; and the broad metabolic and transcriptional rescue achieved by nicotinamide riboside identifies NAD^+^ repletion as the most accessible therapeutic entry point into this axis. Together, these findings establish a mechanistic and pharmacologically tractable framework for brown adipose dysfunction in mitochondrial disease.

## Figure Legends

**Supp. Fig. 1. mtDNA deletion load and mitochondrial Complex II protein abundance in Deletor BAT.**

(A) Long-range PCR analysis of mitochondrial DNA deletion products in interscapular BAT from Deletor mice. Multiple mtDNA deletion products of variable size are detected in all Deletor samples, since WT mice show no detectable mtDNA deletions, confirming the presence of mitochondrial genome instability in this tissue. The short-range PCR product serves as an internal control for total mtDNA content.

(B) Representative immunofluorescence images of BAT sections from wild type (WT) and Deletor (Del) mice stained for DAPI (blue) and SDHA (succinate dehydrogenase subunit A, Complex II-red). Scale bar = 40μm.

(C) Quantification of SDHA staining intensity per nucleus in WT and Deletor BAT (n=5 (WT) n=3 (Del)).

(D) Whole-body oxygen consumption (vO2, ml/kg/hr) and respiratory exchange ratio (RER, vCO2/vO2) measurements by indirect calorimetry (CLAMS; Columbus Instruments) in WT and Deletor mice across light, dark and cold exposure (+4C) phases (n=5 (WT), n=6 (Del)). Top panels show continuous measurements over the 28-hour period and bottom panels show quantification of vO2 and RER within light, dark and cold phases.

Data are shown as mean ± SEM with individual data points overlaid. Statistical analysis was performed by unpaired t-test. For image quantification, 5 images were acquired from each animal and average used for the analysis.

**Supp. Fig. 2. MCT1-defined adipocyte morphology is not altered by pharmacological modulation of the PHD3–HIF axis or iron loading.**

(A) MCT1 immunostaining and quantification of cellular volume per nucleus in WT and Deletor BAT from vehicle-, PX-478-, roxadustat-, and iron dextran-treated mice. MCT1-defined cellular volume per nucleus was not significantly altered by any treatment in either genotype, indicating that adipocyte atrophy in Deletor BAT is refractory to pharmacological perturbation of the pseudohypoxic program. Scale bar = 30 μm.

(B) Representative 3D reconstructions of vascular networks generated using Imaris software. Scale bar = 40μm.

**Supp. Fig. 3. Thermogenic identity is not restored by HIF axis modulation and is further suppressed by iron loading.**

(A) Representative immunofluorescence images of BAT sections stained for DAPI and UCP1 from vehicle-, PX-478-, roxadustat-, and iron dextran-treated wild-type and Deletor mice. UCP1 immunoreactivity remained markedly reduced in all Deletor treatment arms relative to wild-type controls and was further suppressed by iron dextran treatment, demonstrating that thermogenic identity is not downstream of HIF signalling alone and that iron loading independently compounds thermogenic failure. Scale bar = 20 μm.

(B) Perl’s Prussian Blue staining of BAT sections of WT and Deletor from vehicle-, PX-478-, roxadustat-, and iron dextran-treated mice. Scale bar = 60 µm.

**Supp. Fig. 4. Macrophage accumulation and transcriptional pathway alterations following HIF modulation and iron loading in BAT.**

(A) Representative immunohistology images of BAT sections stained for CD68 from vehicle-, PX-478-, roxadustat-, and iron dextran-treated WT and Deletor mice. Scale bar = 60 μm.

(B) KEGG pathway enrichment analysis of differentially expressed genes in WT treatment relative to WT vehicle BAT following PX-478, roxadustat, and iron dextran treatment, showing the top significantly enriched upregulated and downregulated pathways. Pathways are ranked according to enrichment significance and gene representation within each pathway.

**Supp. Fig. 5. Metabolomic profiling of BAT following nicotinamide riboside treatment**

(A) Heatmap showing treatment-dependent regulation of the most significantly altered metabolites across WT, WT-NR, Deletor, and Deletor-NR groups. Relative metabolite abundance is represented by row-wise normalized intensity values. (n=6 per group)

(B) Metabolite pathway enrichment analysis of NR-responsive metabolites in WT BAT, highlighting purine metabolism and glycerophospholipid metabolism as the most significantly enriched pathways. Bubble size indicates the number of pathway hits, and colour intensity indicates statistical significance.

(C) Volcano plot of differentially abundant metabolites in WT-NR versus WT BAT, with selected significantly altered metabolites highlighted. Metabolomic analyses were performed using BAT samples from WT and WT-NR (n = 6 per group) mice.

(D) Expression of extracellular matrix remodelling genes in BAT from Del vehicle and Del NR mice, including *Col1a1, Col1a2, Col3a1, Mmp14, Ddr2, Thbs1, Fn1, Lama2,* and *Tnxb* (Del vehicle and Del NR (n = 4), WT vehicle and WT NR (n=5).

(E) Expression of thermogenesis-associated genes in BAT from Del vehicle and Del NR mice, including *Cycs, Ucp1, Fabp3, Prdm16, Ppargc1a, Cpt1b, Cox7a1, Cox8b, Dio2,* and *Cidea* (Del vehicle and Del NR (n=4), WT vehicle and WT NR (n=5)

(F) Relative abundance of free carnitine and selected acylcarnitine species in WT vehicle, WT-NR, Deletor vehicle, and Deletor-NR BAT, showing restoration of carnitine-associated metabolites in Deletor BAT after NR treatment. (n=6 per group)

Data are shown as mean ± SEM. Statistical significance was assessed by one-way ANOVA followed by Tukey’s multiple comparisons test.

## Methodology

### Mouse model

This study utilized the Deletor mouse model, which carries a dominant *Twnk* gene mutation (duplication of amino acids 353–365) encoding the mitochondrial DNA helicase Twinkle, on a C57BL/6 congenic background [7]. Age-matched wild-type (WT) littermates were used as controls. All experiments were performed in 24-month-old male twinkle-D line. Mice were housed individually under controlled conditions (22°C, 12 h light/dark cycle) with *ad libitum* access to standard rodent chow (Altromin Spezialfutter GmbH & Co. Cat. No. 1324, KG; 11% fat, 65% carbohydrate, 24% protein by caloric content). Body weight and food intake were monitored regularly. All animal procedures were conducted in accordance with ethical guidelines and approved by the State Provincial Office for Animal Experimentation of Finland.

### Treatments

The transgenic Deletor mice were treated as following:

#### PX-478 (HIF inhibitor)

Mice were administered PX-478 (25 mg/kg) (MedChemExpress, HY-10231) by intraperitoneal (i.p.) injection three times per week for 6 weeks. PX-478 was dissolved in a vehicle containing 5% DMSO, 40% PEG300, 5% Tween-80, and 50% saline. Vehicle-treated mice received equivalent volumes of the vehicle.

#### Roxadustat (HIF stabilizer)

Mice were administered roxadustat (50 mg/kg) (MedChemExpress, HY-13426) by i.p. injections three times per week for 6 weeks. Roxadustat was dissolved in a vehicle consisting of 5% DMSO, 40% PEG300, 5% Tween-80, and 50% saline. Vehicle-treated mice received equivalent volumes of the vehicle.

#### Iron Dextran

Mice were administered iron dextran (FeD) (60mg/kg) (Sigma-Aldrich, #D8517) by i.p. injections three times a week. Iron Dextran was diluted in saline. Vehicle-treated controls received equivalent volumes of dextran (Dex) with saline to introduce acute iron stress.

#### Nicotinamide Riboside (NR)

Mice received NR (400 mg/kg in chow, Chromadex) for six months as described previously [28]

### Respiratory Function Testing in Animals

Oxygen consumption and carbon dioxide production were measured in Oxymax Lab Animal Monitoring System (CLAMS; Columbus Instruments). In CLAMS the mice were housed in individual cages in temperature-regulated chambers, settled for 24 hours and were recorded for 28 hours. The settling and the first day of recording were in room temperature (+22°C) and for 6 hours of acute cold exposure at (+4°C). Respiratory exchange ratio (RER) was calculated to indicate a rough estimate of the preferred fuel (carbohydrate breakdown vs. fatty acid oxidation) of the organism.

### Tissue collection and processing

Brown adipose tissue (BAT) was collected immediately after sacrifice and fixed in 10% neutral-buffered formalin, then dehydrated through graded alcohols and xylene and embedded in paraffin (FFPE). For Oil Red O lipid staining analyses, BAT was embedded in OCT compound (Tissue-Tek) and snap-frozen in 2-methylbutane pre-cooled with liquid nitrogen.

### Immunofluorescence staining

FFPE BAT sections (10 μm) were deparaffinized in xylene or Tissue-Tek® Tissue-Clear® Xylene Substitute (Tissue-Tek, 1466) (3 × 5–7 min) and rehydrated through a descending ethanol series. Heat-induced antigen retrieval was performed in Tris–EDTA buffer (pH 9.0) by microwave heating, followed by cooling and 2 × PBS washes. Sections were blocked for 1 h at room temperature in blocking buffer (0.2% BSA and 2.5% normal donkey serum in PBS). Primary antibodies diluted in blocking buffer were applied overnight at 4°C in a humidified chamber: anti-podocalyxin (R&D Systems, AF1556-SP, 1:400), anti-MCT1 (Merck/MilliporeSigma, AB1286-I, 1:500), anti-SDHA (Abcam, ab14715, 1:100), anti-PHD3 (Novus Biologicals, NB100-139SS, 1:250), anti-UCP1 (Abcam, ab10983, 1:500), and anti-FTH1 (Abcam, ab287968, 1:200). Prior to SDHA immunostaining, sections were incubated with M.O.M. (Mouse on Mouse) Blocking Reagent following the manufacturer’s protocol. Following washes in PBS, or Tris–NaCl–Tween 20 (TNT) buffer for stainings with MCT1 and podocalyxin, sections were incubated for 1 h at room temperature in the dark with the appropriate fluorophore-conjugated secondary antibodies diluted 1:500 in 0.05% Tween 20 in PBS. Secondary antibodies included chicken anti-mouse Alexa Fluor 647 (Invitrogen/Molecular Probes, A21463), donkey anti-rabbit IgG H&L Alexa Fluor 594 (Abcam, ab150068), chicken anti-goat Alexa Fluor 594 (Molecular Probes, A21468), donkey anti-chicken Alexa Fluor 488 (Jackson ImmunoResearch, 703-546-155), and donkey anti-goat Alexa Fluor 594 (Invitrogen, A32758). Nuclei were counterstained with DAPI (1:100). After secondary antibody incubation, sections were washed in TNT buffer and PBS, post-fixed in 1% paraformaldehyde for 5 min, washed 2 × PBS and 1 × mQ-H₂O, and mounted in ProLong™ Gold Antifade Mountant (Invitrogen, P36930). Images were acquired using laser-scanning confocal microscopes (LSM880, Zeiss; Stellaris 8 FALCON/DLS, Leica).

### Immunohistochemistry

For CD68 staining, 10 μm FFPE BAT sections were stained as follows. Following deparaffinization, rehydration, and antigen retrieval in Tris–EDTA buffer (pH 9.0), endogenous peroxidase activity was quenched with 0.3% H₂O₂ for 6 min. Sections were blocked in 7.6% normal horse serum in 0.1% Tween-20 in PBS for 1 h and incubated overnight at 4°C with anti-CD68 antibody (1:600). After washing, a polymer-based secondary antibody (ImmPRESS Goat Anti-Rat IgG Polymer Detection Kit, ready-to-use; Vector Laboratories). was applied for 30 min, and the signal was developed with 3,3′-diaminobenzidine (DAB, ∼2 min). Nuclei were counterstained with hematoxylin. Sections were dehydrated, cleared in xylene, and mounted. Imaging was done by light microscopy (Axioplan 2 Universal Microscope, Zeiss)

### Histochemical staining

#### Picrosirius Red

Collagen deposition was assessed on FFPE sections (8-10 μm). Following deparaffinization and rehydration, sections were incubated in Sirius Red solution for 20 min, briefly differentiated in acetic acid, dehydrated through graded ethanol, cleared in xylene, and mounted for brightfield and polarized light microscopy (Axioplan 2 Universal Microscope, Zeiss).

#### Perl’s Prussian Blue

Iron accumulation was assessed on FFPE BAT sections (10 μm). After deparaffinization and rehydration, sections were incubated for 10–20 min in freshly prepared 5% potassium ferrocyanide/5% hydrochloric acid (or 5% potassium ferrocyanide in 10% hydrochloric acid for aged-mouse cohorts). Sections were washed thoroughly, counterstained with Nuclear Fast Red (0.1%, 3 min; Vector Laboratories), dehydrated, cleared in xylene, and mounted in Pertex (Histolab, Z00811). Liver sections served as positive control.

#### Oil Red O

Lipid content was assessed on OCT-embedded frozen sections (8–10 μm). Sections were fixed in formic calcium solution (concentrated formalin: water:10% CaCl₂, 1:10:1) and incubated for 10 min in freshly prepared Oil Red O working solution. The working solution was prepared from a commercially available Oil Red O stock solution (Sigma-Aldrich, O1391-500ML) by dilution with water and filtration immediately before use. Nuclei were counterstained with Mayer’s hematoxylin. Sections were mounted in Aquatex aqueous mounting medium (Merck, 1.02419) and imaged by light microscopy (Axioplan 2 Universal Microscope, Zeiss).

### Image analysis

#### Oil Red O quantification

Quantitative analysis of Oil Red O-stained sections was performed using a custom batch-processing macro in FIJI/ImageJ (v2.16.0). Images acquired at 40× magnification (3072 × 2300 pixels) were split into RGB channels, and the blue channel was used for analysis due to optimal contrast. A fixed grey-level threshold of 0–136, determined empirically from representative images, was applied uniformly across all samples. Thresholded images were converted to binary masks and analysed using the “Analyse Particles” function; objects smaller than 800 μm² or intersecting image borders were excluded with no circularity restriction. Particle count, mean particle area, and total detected area were exported for downstream statistical analysis.

#### Vascular morphometry

BAT vasculature was quantified from podocalyxin-immunolabelled FFPE sections imaged as Z-stacks in .czi format using a confocal microscope. Images were opened in FIJI/ImageJ (v2.16.0) through Bio-Formats; where a maximum intensity projection was generated from z-stacks. FIJI is an open-source distribution of ImageJ for biological image analysis [59]. Images were converted to 8-bit and subjected to rolling ball background subtraction (radius = 50 pixels). Images were inverted prior to thresholding. Binary masks were generated by automatic Otsu thresholding and skeletonised using the Skeletonize command. Vascular topology was quantified with Analyse Skeleton (2D/3D), yielding per-image metrics including branch count, junction count, mean and maximum branch length, and the longest shortest path within the vascular graph. Voxel size metadata extracted from Bio-Formats were used to convert all length measurements from voxels to micrometres. For volumetric analysis, Z-stack images were imported into Imaris (Bitplane); three-dimensional surface reconstructions were generated from the podocalyxin channel, and DAPI-labelled nuclei were detected as discrete objects. Vascular volume was normalised to nuclear count (vascular area/DAPI) to account for differences in tissue size, volume, and section thickness across samples.

### mtDNA deletion load analysis

Total DNA was extracted from snap-frozen BAT by phenol-chloroform extraction followed by ethanol precipitation. mtDNA deletion load was assessed by long-range and short-range PCR using Phusion High-Fidelity DNA Polymerase with GC buffer (Thermo Fisher Scientific, F-530).

Long-range PCR was performed to detect deletion products spanning the major arc using the following primers: 5′-GGTTCGTTTGTTCAACGATTAAAGTCCTACGTG-3′ (forward) and 5′-GAGGTGATGTTTTTGGTAAACAGGCGGGGT-3′ (reverse). Each 50 μL reaction contained 1× GC buffer, 0.5 μM of each primer, 200 μM dNTPs, 0.5 U Phusion polymerase, and 10 ng template DNA. Cycling conditions were as follows: 98°C for 30 s; 22 cycles of 98°C for 10 s and 72°C for 3 min; final extension at 72°C for 10 min.

Short-range PCR targeting a non-deleted mtDNA region (549 bp) was performed as an internal control for mtDNA content using the primers: 5′-ACCCCGCCTGTTACCAAAAACATCACCTC-3′ (forward) and 5′-ACGTACCACTTTAATCGTTGAACAAACGAACC-3′ (reverse). Each 15.7 μL reaction contained 1× GC buffer, 0.83 μM of each primer, 318 μM dNTPs, 0.2 U Phusion polymerase, and 10 ng template DNA. Cycling conditions were: 98°C for 30 s; 20 cycles of 98°C for 10 s and 72°C for 2 min; final extension at 72°C for 10 min. PCR products were resolved on 1% agarose gels and visualised using a ChemiDoc imaging system (Bio-Rad).

### Electron microscopy

For ultrastructural analysis, brown adipose tissue (BAT) samples were fixed in 2.5% glutaraldehyde prepared in 0.1 M phosphate buffer, followed by post-fixation in 1% osmium tetroxide. Samples were dehydrated through a graded ethanol series and embedded in epoxy resin. Semi-thin sections (1 μm) were stained with 0.5% toluidine blue and 1% boric acid and examined by light microscopy to identify regions of interest. Ultrathin sections (60–90 nm) were then cut from selected regions, mounted on copper grids, and contrasted with uranyl acetate and lead citrate. Images were acquired using a JEOL JEM-1400 transmission electron microscope.

### RNA isolation and cDNA synthesis

Total RNA was extracted from snap-frozen BAT using TRIzol reagent (Invitrogen) with tissue homogenisation performed in a Precellys 24 homogenizer (Bertin Technologies). RNA was further purified using the NucleoSpin RNA Clean-up XS kit (Macherey-Nagel, 740903.5) according to the manufacturer’s instructions, including on-column rDNase digestion (Macherey-Nagel, 740963) to eliminate genomic DNA contamination. RNA concentration and integrity were assessed using an Agilent TapeStation system. Samples with a minimum of RIN value 7 were used for RNA-sequencing.

### Targeted Metabolomics Data Analysis

Metabolites were extracted from brown adipose tissue sections around 10 mg by adding 500 μL of cold extraction solvent (acetonitrile/methanol/Milli-Q water 40:40:20, Thermo Fisher Scientific) followed by homogenization using a tissue homogenizer (Bertin Technologies) for three cycles (30 s at 5500 rpm in +4°C with 60 s pauses in between cycles). The homogenized samples were centrifuged at 14 000 rpm in +4°C for 5 min, after which 450 μL of sample supernatants were filtered through Phree Phospholipid Removal plates (Phenomenex). 300 μL of filtrated samples were transferred into polypropylene tubes and evaporated dry under nitrogen stream. Dried samples were reconstituted in 40 μL of extraction solvent and vortexed for 2 min before being transferred into high-performance liquid chromatography glass autosampler vials. Samples were analyzed using a Thermo Vanquish UHPLC coupled with Q-Exactive Orbitrap quadrupole mass spectrometer using MS1 scanning mode. The data quality was monitored throughout the run using a pooled quality control (QC) sample, which had been prepared by combining an aliquot of 2 μL from each reconstituted sample. Injections of the QC sample were interspersed throughout the run after every ten samples. The data processing and integration of metabolites was performed with TraceFinder 5.2 software (Thermo Fisher Scientific) using confirmed retention times from in-house standard library (MSMLS-1EA, Merck) and their *m/z* (5 ppm). The metabolite data was quality controlled for peak quality (poor chromatographs), relative standard deviation (with 20% cutoff) and carry-over (with 20% cutoff). Data was normalized to the total sum intensity (peak areas).

### Transcriptomic Analysis

Total RNA was isolated from WT (n = 5) and Deletor (n =4) mice per treatment group and subjected to transcriptome profiling. RNA was extracted from snap-frozen brown adipose tissue (BAT) samples using TRIzol reagent (Invitrogen). Tissue homogenization was performed with the Precellys Lysing Kit – Tissue Homogenizing CKMix (Bertin Technologies) using the Precellys 24 homogenizer (Bertin Technologies). The extracted RNA was treated with DNase and purified using RNeasy Mini columns (Qiagen) according to the manufacturers’ instructions. RNA integrity was assessed using the TapeStation system, and only samples with an RNA Integrity Number (RIN) ≥ 7 were used for RNA sequencing.

Library preparation and high-throughput sequencing of WT and Del BAT samples were performed at Beijing Genomics Institute (BGI, Shenzhen, China) according to the manufacturer’s standard protocols. Sequencing was carried out on the BGISEQ-500 platform using 100 bp paired-end (PE100) reads. Differential gene expression analysis was performed in the R statistical environment using the DESeq2 package with default parameters.

For the PX-478-,roxadustat-, and iron dextran treated samples (n=5 per group), 3′ bulk RNA sequencing was performed by Scellex Oy (Helsinki, Finland). During library preparation, 3′-terminal cDNA fragments were selectively enriched by index PCR using Nextera-compatible Illumina index primers together with custom primers complementary to the PCR handle. The resulting libraries were sequenced according to the provider’s standard workflow, and differential gene expression analysis was performed using the DESeq2 package in R with default parameters.

### Quantification and statistical analysis

Histological image quantification was performed in FIJI/ImageJ (v2.16.0, NIH) and Imaris (Bitplane) as described above. Statistical analyses were performed using GraphPad Prism (v10.5.0, GraphPad Software).

Metabolomics data were preprocessed in MetaboAnalyst 6.0 (Xia *et al*, 2009), comprising feature filtering by 20% interquartile range (IQR), intensity sum normalisation, log transformation, and autoscaling. Multiple testing correction was applied using the Benjamini–Hochberg method (false discovery rate threshold: 0.2). Statistical significance for pairwise and multi-group comparisons was assessed by unpaired Student’s *t*-test or one-way ANOVA, as indicated in the figure legends.

Transcriptomics data were evaluated by principal component analysis (PCA) as an unsupervised approach to assess sample clustering and group separation. Pathway enrichment analysis was performed using ShinyGO (v0.85; Ge *et al*, 2020). Statistical significance for pairwise and multi-group comparisons was assessed by unpaired Student’s *t*-test or one-way ANOVA, as indicated in the figure legends.

## Data availability

The RNA-sequencing data generated in this study are available in the European Nucleotide Archive (ENA) under accession number PRJEB115293. Raw metabolomics mass spectrometry data have been deposited in the Mass Spectrometry Interactive Virtual Environment (MassIVE) repository and can be accessed under accession number MSV000102227.

## Acknowledgements

We thank the staff of the Laboratory Animal Centre at the University of Helsinki for expert animal husbandry and veterinary support. We are grateful to Professor Anu Suomalainen (University of Helsinki) for generously providing the Deletor mice used in this study. Nicotinamide riboside was kindly provided by Niagen (previously Chromadex) for use in this study. We thank Helena Puro for technical assistance and dedicated support throughout the experimental work.

Metabolomic analyses were performed at the Metabolomics Core Facility, UT Southwestern Medical Center, Dallas, Texas and we are grateful to the facility staff for technical assistance and data processing support. Transcriptomic sequencing was performed by the FUGU Sequencing Unit at the Helsinki Institute of Life Science (HiLIFE) and Biocenter Finland, University of Helsinki, and we thank the unit staff for library preparation, sequencing, and primary data processing. Transcriptomic sequencing and library preparation for hypoxia and iron treatment samples were performed at Scellex Oy, Biomedicum, Helsinki, Finland, and we thank the facility staff for technical assistance and data processing. Transcriptomics sequencing and library preparation for NR treatment was performed at BGI, China, and we thank the facility staff for data processing. Electron microscopy and confocal imaging were performed at the Electron Microscopy Unit and the Light Microscopy Unit of the Helsinki Institute of Life Science (HiLIFE), University of Helsinki, and we thank the facility staff for expert assistance with sample preparation and image acquisition. The authors acknowledge the Biomedicum Imaging Unit (BIU), University of Helsinki, for providing access to microscopy facilities in fluorescent imaging and technical support. Madeleine H Lackman was supported by the Doctoral Programme in Integrative Life Science, University of Helsinki, the Biomedicum Helsinki Foundation, and the Orion Research Foundation. Sinem Karaman was supported by the Research Council of Finland (grants #330053, #358080, and #360042) and the Sigrid Jusélius Foundation. The Children’s Research Institute Metabolomics Facility is supported by the Cancer Prevention Research Institute of Texas (CPRIT Core Facilities Support Award RP240494). This work was supported by the Research Council of Finland, Nahid A Khan (grants #316435 and #355637).

