## Supplemental Information for "Mitochondrial genome instability disrupts brown adipose tissue through pseudohypoxia-iron-NAD⁺ axis"

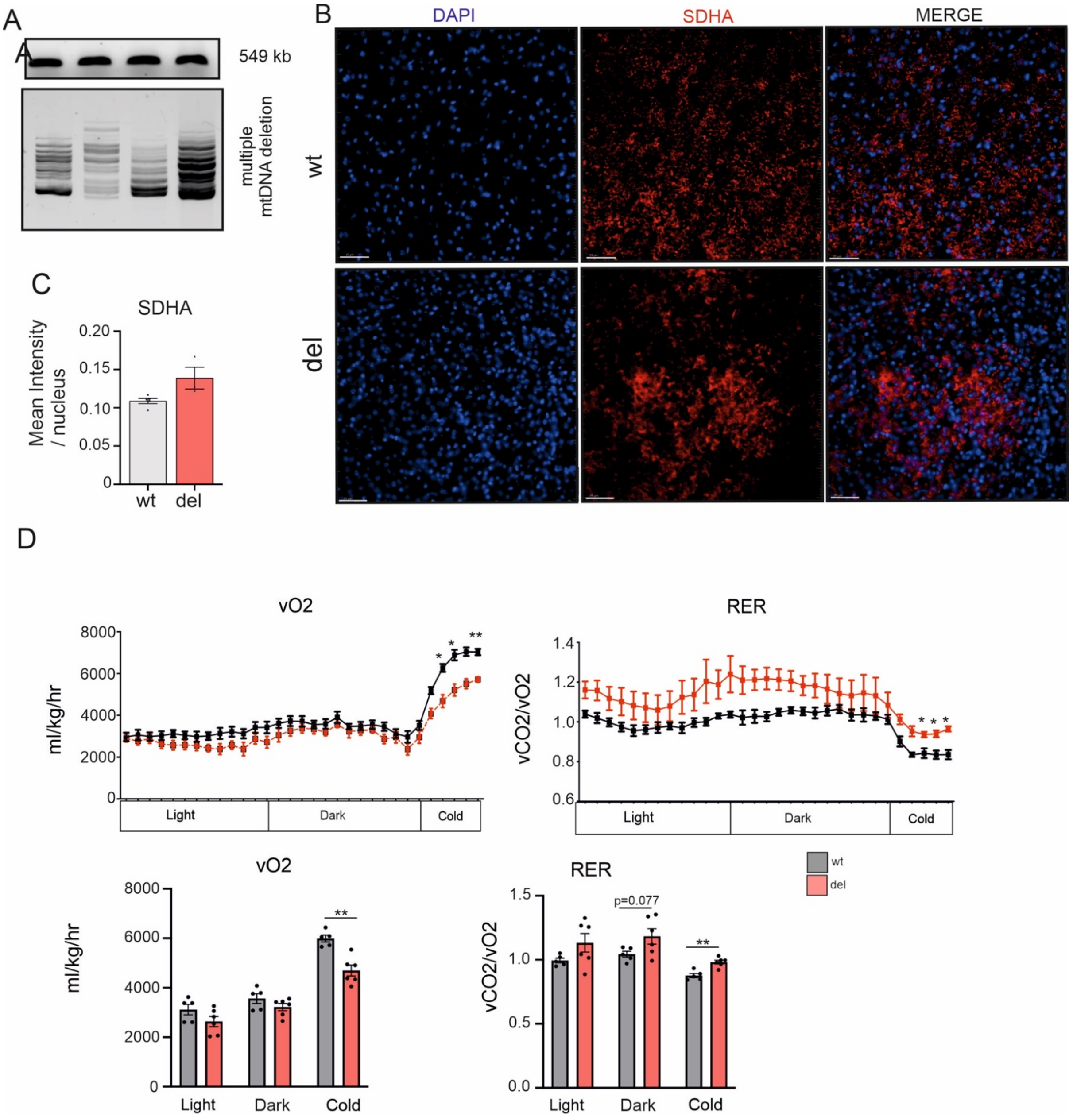


**Supp. Fig. 1. mtDNA deletion load and cold-induced thermogenic failure in Deletor BAT.**

(A) Long-range PCR analysis of mitochondrial DNA deletion products in interscapular BAT from Deletor mice. Multiple mtDNA deletion products of variable size are detected in all Deletor samples, since WT mice show no detectable mtDNA deletions, confirming the presence of mitochondrial genome instability in this tissue. The short-range PCR product serves as an internal control for total mtDNA content.

(B) Representative immunofluorescence images of BAT sections from wild type (WT) and Deletor (Del) mice stained for DAPI (blue) and SDHA (succinate dehydrogenase subunit A, Complex II-red). Scale bar = 40μm.

(C) Quantification of SDHA staining intensity per nucleus in WT and Deletor BAT (n=5 (WT) n=3 (Del)).

(D) Whole-body oxygen consumption (vO2, ml/kg/hr) and respiratory exchange ratio (RER, vCO2/vO2) measurements by indirect calorimetry (CLAMS; Columbus Instruments) in WT and Deletor mice across light, dark and cold exposure (+4C) phases (n=5 (WT), n=6 (Del)). Top panels show continuous measurements over the 28-hour period and bottom panels show quantification of vO2 and RER within light, dark and cold phases.

Data are shown as mean ± SEM with individual data points overlaid. Statistical analysis was performed by unpaired t-test. For image quantification, 5 images were acquired from each animal and average used for the analysis.

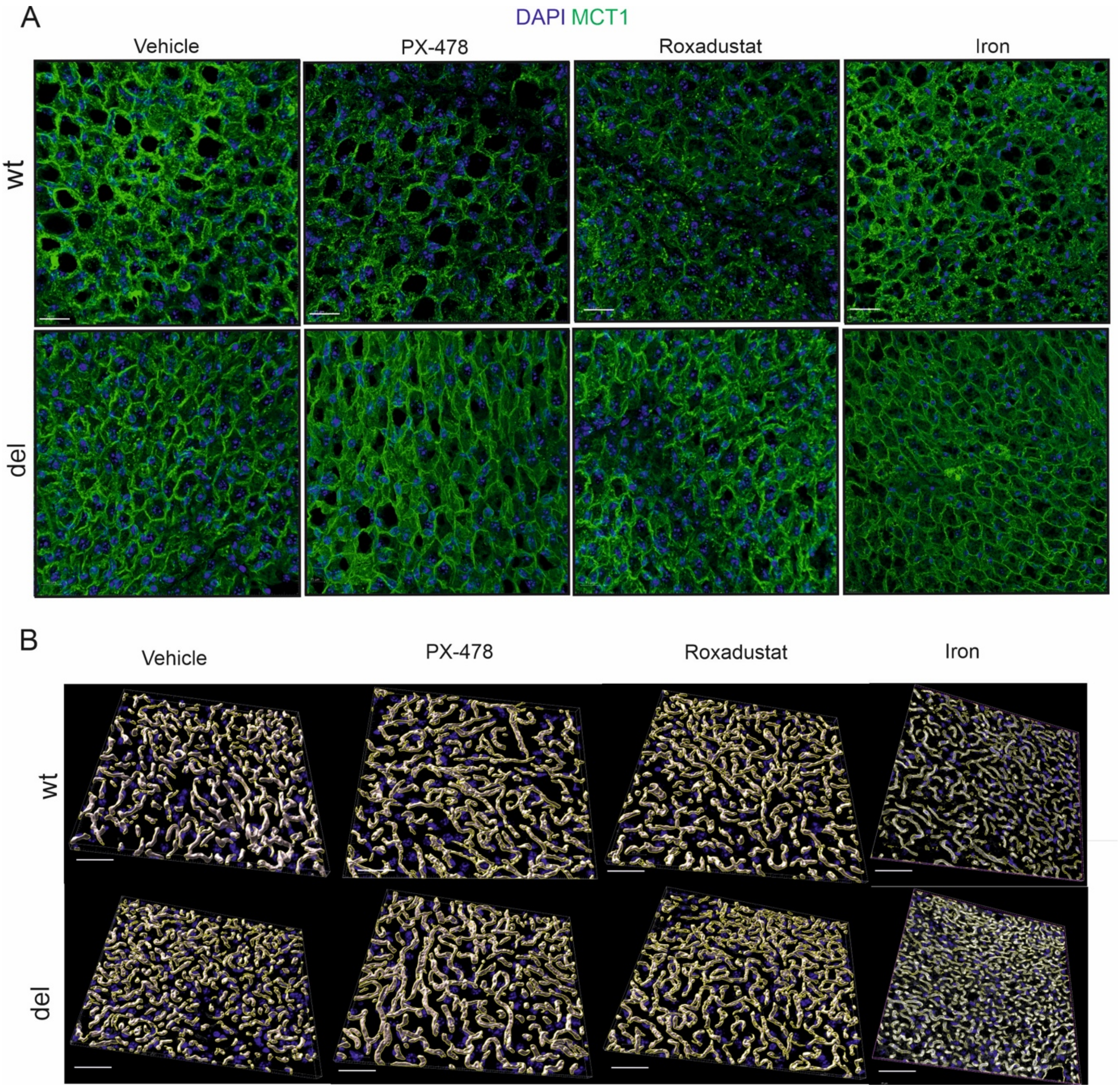


**Supp. Fig. 2. MCT1-defined adipocyte morphology is not altered by pharmacological modulation of the PHD3–HIF axis or iron loading.**

(A) MCT1 immunostaining and quantification of cellular volume per nucleus in WT and Deletor BAT from vehicle-, PX-478-, roxadustat-, and iron dextran-treated mice. MCT1-defined cellular volume per nucleus was not significantly altered by any treatment in either genotype, indicating that adipocyte atrophy in Deletor BAT is refractory to pharmacological perturbation of the pseudohypoxic program. Scale bar = 30 μm.

(B) Representative 3D reconstructions of vascular networks generated using Imaris software. Scale bar = 40μm.

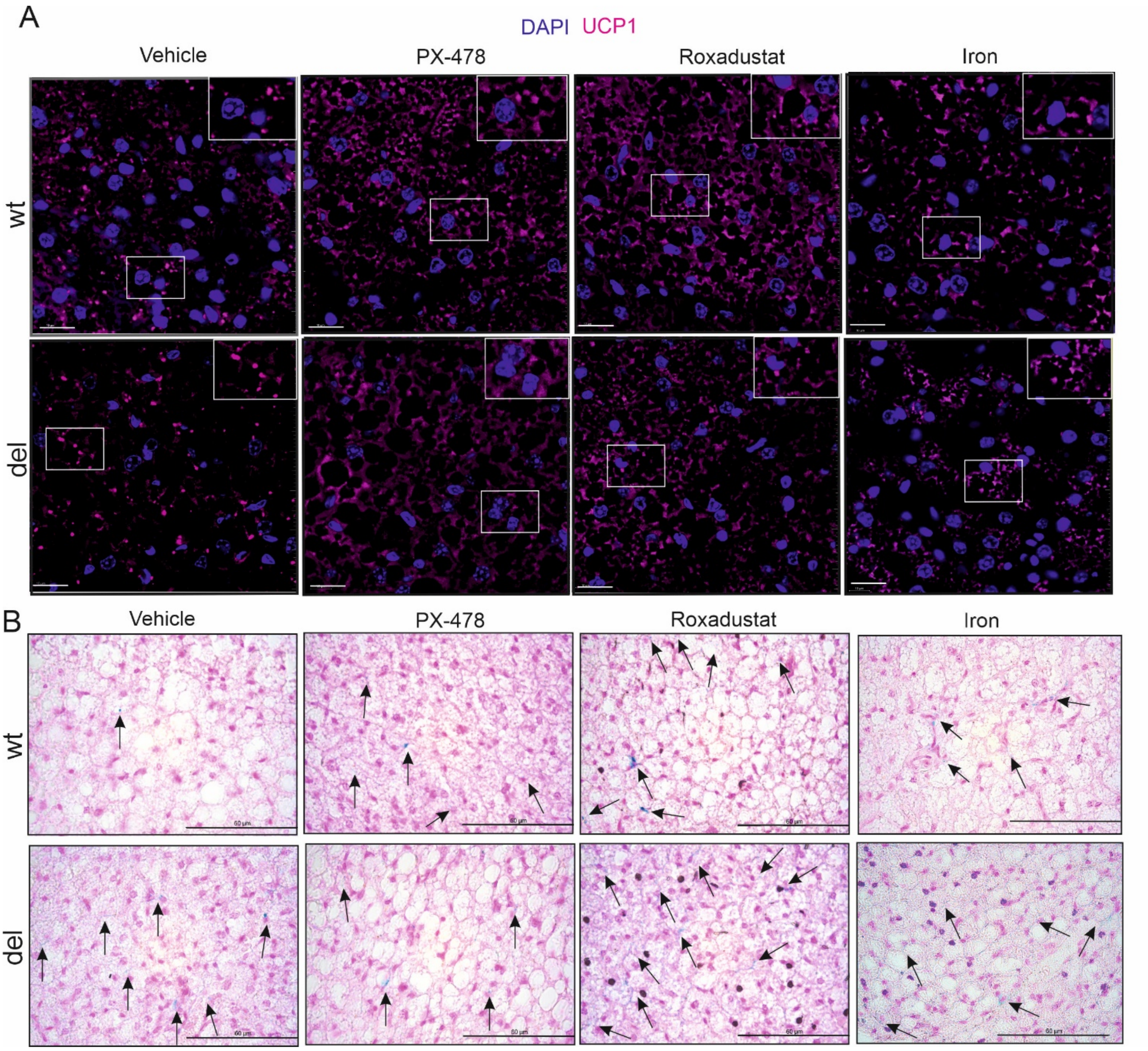


**Supp. Fig. 3. Thermogenic identity is not restored by HIF axis modulation and is further suppressed by iron loading.**

(A) Representative immunofluorescence images of BAT sections stained for DAPI and UCP1 from vehicle-, PX-478-, roxadustat-, and iron dextran-treated wild-type and Deletor mice. UCP1 immunoreactivity remained markedly reduced in all Deletor treatment arms relative to wild-type controls and was further suppressed by iron dextran treatment, demonstrating that thermogenic identity is not downstream of HIF signalling alone and that iron loading independently compounds thermogenic failure. Scale bar = 20 µm.

(B) Perl's Prussian Blue staining of BAT sections of WT and Deletor from vehicle-, PX-478-, roxadustat-, and iron dextran-treated mice. Scale bar = 60 μm.

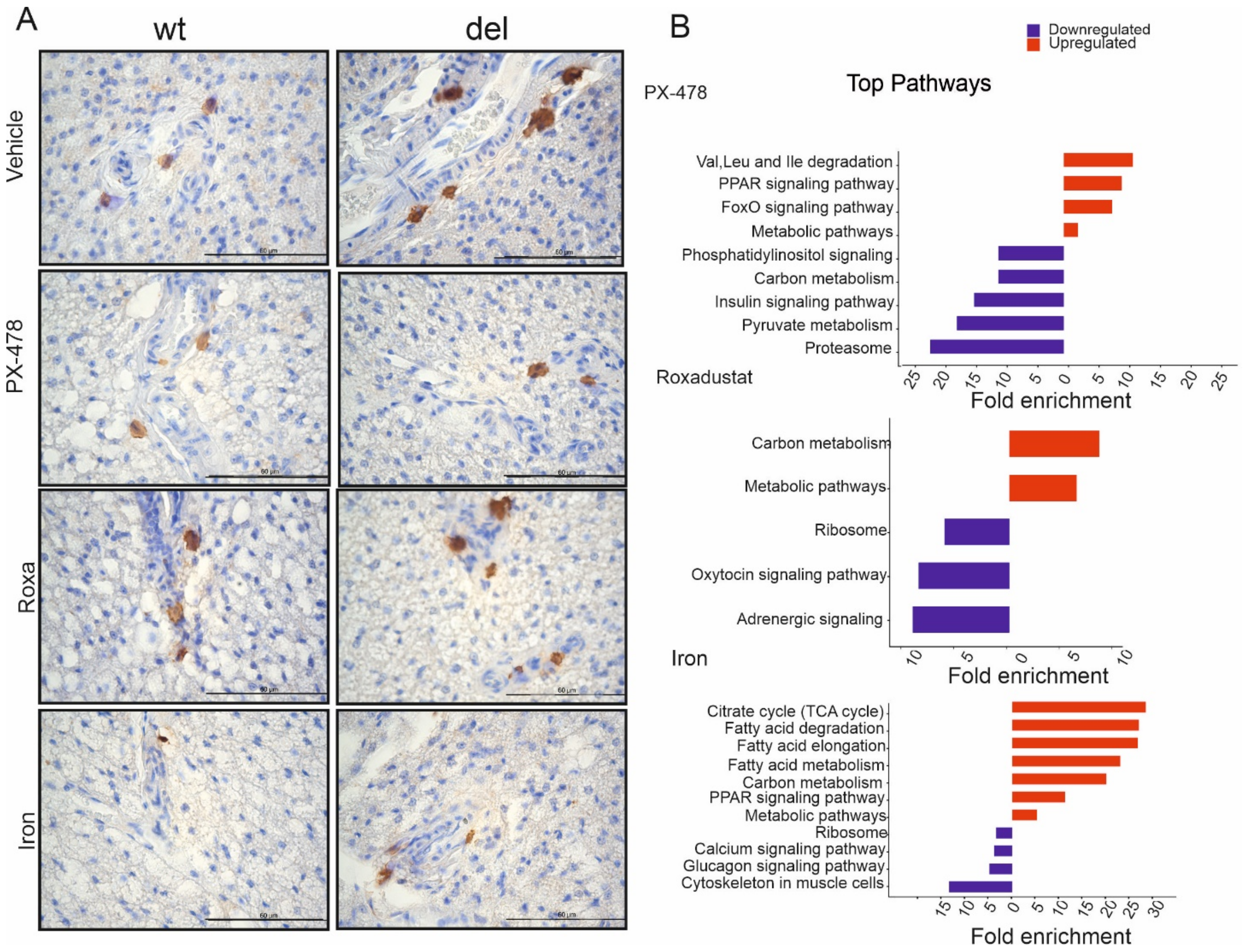


**Supp. Fig. 4. Macrophage accumulation and transcriptional pathway alterations following HIF modulation and iron loading in BAT.**

(A) Representative immunohistology images of BAT sections stained for CD68 from vehicle-, PX-478-, roxadustat-, and iron dextran-treated WT and Deletor mice. Scale bar = 60 μm.

(B) KEGG pathway enrichment analysis of differentially expressed genes in WT treatment relative to WT vehicle BAT following PX-478, roxadustat, and iron dextran treatment, showing the top significantly enriched upregulated and downregulated pathways. Pathways are ranked according to enrichment significance and gene representation within each pathway.

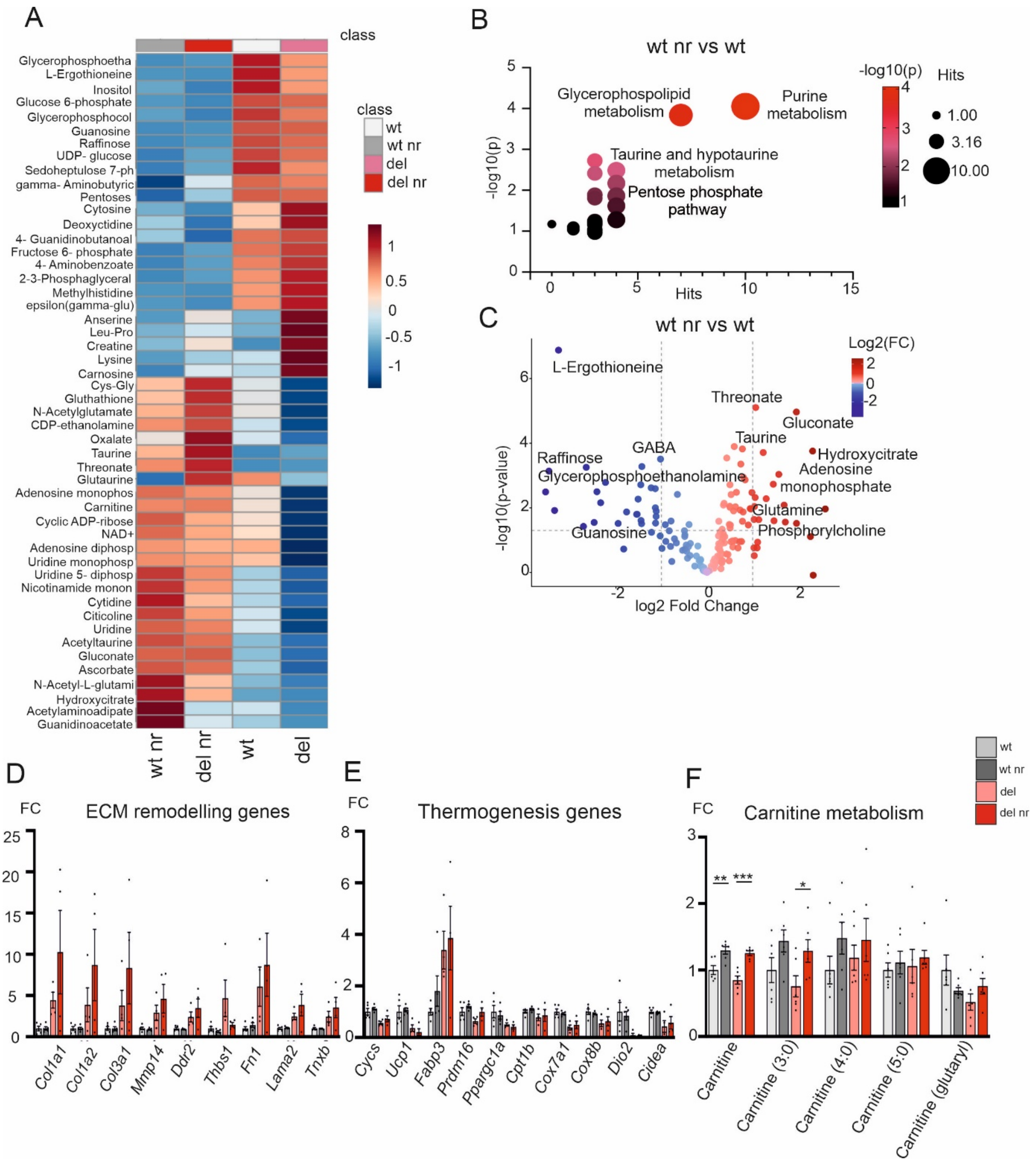


**Supp. Fig. 5. Metabolomic profiling of BAT following nicotinamide riboside treatment**

(A) Heatmap showing treatment-dependent regulation of the most significantly altered metabolites across WT, WT-NR, Deletor, and Deletor-NR groups. Relative metabolite abundance is represented by row-wise normalized intensity values. (n=6 per group)

(B) Metabolite pathway enrichment analysis of NR-responsive metabolites in WT BAT, highlighting purine metabolism and glycerophospholipid metabolism as the most significantly enriched pathways. Bubble size indicates the number of pathway hits, and colour intensity indicates statistical significance.

(C) Volcano plot of differentially abundant metabolites in WT-NR versus WT BAT, with selected significantly altered metabolites highlighted. Metabolomic analyses were performed using BAT samples from WT and WT-NR (n = 6 per group) mice.

(D) Expression of extracellular matrix remodelling genes in BAT from Del vehicle and Del NR mice, including *Col1a1, Col1a2, Col3a1, Mmp14, Ddr2, Thbs1, Fn1, Lama2,* and *Tnxb* (Del vehicle and Del NR (n = 4), WT vehicle and WT NR (n=5).

(E) Expression of thermogenesis-associated genes in BAT from Del vehicle and Del NR mice, including *Cycs, Ucp1, Fabp3, Prdm16, Ppargc1a, Cpt1b, Cox7a1, Cox8b, Dio2,* and *Cidea* (Del vehicle and Del NR (n=4), WT vehicle and WT NR (n=5)
(F) Relative abundance of free carnitine and selected acylcarnitine species in WT vehicle, WT-NR, Deletor vehicle, and Deletor-NR BAT, showing restoration of carnitine-associated metabolites in Deletor BAT after NR treatment. (n=6 per group)

Data are shown as mean ± SEM. Statistical significance was assessed by one-way ANOVA followed by Tukey's multiple comparisons test.
